# Reconstructing Squamate Biogeography in Afro-Arabia Reveals the Influence of a Complex and Dynamic Geologic Past

**DOI:** 10.1101/2020.12.01.406546

**Authors:** Héctor Tejero-Cicuéndez, Austin H. Patton, Daniel S. Caetano, Jiří Šmíd, Luke J. Harmon, Salvador Carranza

## Abstract

The geographic distribution of biodiversity is central to understanding evolutionary biology. Paleogeographic and paleoclimatic histories often help to explain how biogeographic patterns unfold through time. However, such patterns are also influenced by a variety of other factors, such as lineage diversification, that may affect the probability of certain types of biogeographic events. The complex and well-known geologic and climatic history of Afro-Arabia, together with the extensive research on reptile systematics in the region, makes Afro-Arabian squamate communities an ideal system to investigate biogeographic patterns and their drivers. Here we reconstruct the phylogenetic relationships and the ancestral geographic distributions of several Afro-Arabian reptile clades (totaling 430 species) to estimate the number of dispersal, vicariance and range contraction events. We then compare the observed biogeographic history to a distribution of simulated biogeographic events based on the empirical phylogeny and the best-fit model. This allows us to identify periods in the past where the observed biogeographic history was likely shaped by forces beyond the ones included in the model. We find an increase in vicariance following the Oligocene, most likely caused by the fragmentation of the Afro-Arabian plate. In contrast, we did not find differences between observed and expected dispersal and range contraction levels. This is consistent with diversification enhanced by environmental processes and with the establishment of a dispersal corridor connecting Africa, Arabia and Eurasia since the middle Miocene. Finally, here we show that our novel approach is useful to pinpoint events in the evolutionary history of lineages that might reflect external forces not predicted by the underlying biogeographic model.

## INTRODUCTION

One of the central goals of evolutionary biology is to understand why species present certain biogeographic distribution patterns (Cox et al. 1973; Schall and Pianka 1978; Cracraft 1983). Biogeographic studies highlight the role of paleogeography and paleoclimate in shaping the current distribution of diversity (Raven and Axelrod 1972; Cox 1974; Gamble et al. 2011; Giribet et al. 2012; Beaulieu et al. 2013). However, geological and environmental processes may drive not only specific biogeographic histories, but also diversification dynamics (Flagstad et al. 2001; Drovetski 2003; Kozak and Wiens 2006; Ricklefs 2006; Fjeldså et al. 2012; Toussaint et al. 2014; Badgley et al. 2017; Oliveros et al. 2019). In this context, inferring the underlying causes of biogeographic patterns can be challenging, since the incidence of certain biogeographic processes (e.g., dispersal or vicariance) is not only subjected to historical factors, but also to changes in the number of species through time (Sanmartín et al. 2001; Sanmartín and Ronquist 2004).

Some systems on Earth are particularly interesting from a biogeographic perspective due to the nature of their shifting climatic or geologic histories. Such is the case of biotic interchanges, in which tectonic and paleoenvironmental processes have had an impact on the distribution of biodiversity (Marshall et al. 1982; Barry et al. 1991; Vermeij 1991*a*, *b*; Bibi 2011). The region of Afro-Arabia and its biota constitute an ideal system to investigate general biogeographic patterns and their relationship with past environmental conditions and diversification trends. That is, the geologic history of this region is characterized by a complex succession of landmass fragmentation and reconnection events beginning during the Eocene with the split of the Arabian and African plates around 40-30 million years ago (Ma), which eventually triggered the onset of the Red Sea formation around 25 Ma (Bohannon 1986; Ghebreab 1998; Redfield et al. 2003; Bosworth et al. 2005; Scotese 2014a, b). The recurrent formation of dispersal barriers and land bridges, together with constantly shifting climatic conditions (Flower and Kennett 1994; DeMenocal 1995), has potentially left an imprint on the evolutionary history of the biodiversity inhabiting the region (Delany 1989; Winney et al. 2004; Thiv et al. 2010; Šmíd et al. 2013; Metallinou et al. 2015; Tamar et al. 2016*a*, 2018; Simó-Riudalbas et al. 2019). The desert conditions in the Afro-Arabian region contributed to the extensive diversification of arid-adapted taxa, such as reptiles (Brito et al. 2014; Šmíd et al. 2020), making them particularly suitable for studying large-scale biogeographic patterns in this area. However, a comprehensive biogeographic assessment of Afro-Arabian reptiles has yet to be conducted.

Here, we conduct an integrative study of the biogeography and macroevolutionary dynamics of squamates in the Afro-Arabian region. We use statistical models to develop an expectation for the incidence of biogeographic events through time and compare it with observed biogeographic patterns. By definition, all models are simplifications of reality and it is naive to expect any statistical model to fully capture all evidence of biogeographical or diversification events provided by empirical phylogenies. Consequently, we use simulations to exploit model inadequacy, pinpointing time-periods during which the observed evolutionary history of squamates significantly deviates from the expectation based on simplistic models. Such periods might be the result of external forces (e.g., paleogeographic and paleoclimatic histories or ecological interactions) affecting the particular biogeographic history of these animals beyond the origin and extinction of lineages.

We found that although a large portion of the observed biogeographic events can be explained by the fitted model and number of lineages through time, some time-periods were more likely shaped by processes other than diversification dynamics alone, being characterized by model inadequacy. We also show that we may unravel how biogeographic history relates to diversification trends by comparing biogeographic expectations to empirical observations and how both are associated with other historical or ecological processes.

## MATERIALS AND METHODS

### Taxon Sampling and Definition of Groups of Interest

In order to investigate the history of biogeographic events involving Africa and Arabia, we first identified all genera within the order Squamata that are currently present in both areas, and then sampled all species in those genera following the most updated taxonomy (Uetz et al. 2020), including species currently in the process of being described. Once we had a phylogenetic tree (see below), we focused our biogeographic assessment on 21 monophyletic groups: five gekkotan groups represented by the genera *Hemidactylus*, *Pristurus*, *Ptyodactylus*, *Stenodactylus,* and *Tropiocolotes*; two scincoid groups, one represented by the genus *Chalcides* and the other by the clade encompassing the genera *Eumeces*, *Scincopus,* and *Scincus*; two lacertoid groups corresponding to the genera *Acanthodactylus* and *Mesalina*; three iguanian groups, including the genus *Chamaeleo*, the genus *Uromastyx*, and the clade containing the genera *Acanthocercus*, *Pseudotrapelus,* and *Xenagama*; one anguimorph group represented by the genus *Varanus*; and eight snake groups corresponding to the genera *Atractaspis*, *Bitis*, *Cerastes*, *Echis*, *Malpolon*, *Naja*, *Psammophis,* and *Telescopus*. We sampled all the available species from each of these groups except for *Hemidactylus*, for which we considered the “arid clade” only (Carranza and Arnold 2006) as a group of interest for the purpose of this study. This clade is defined by the node including *H. laevis* and *H. granosus* in our phylogenetic reconstruction, and comprises 53 described species. We excluded the Afro-Arabian snake genus *Lytorhynchus* from the biogeographic analysis because we only had two species, precluding any biogeographic reconstruction. We further sampled at least one species representing each family of Squamata and additional species to serve as outgroups and aid in divergence time calibrations, including the sister species to Squamata, the tuatara (*Sphenodon punctatus*), that was used as outgroup. In total, we sampled 555 species: 430 species belonging to the clades of interest and included in the subsequent biogeographic analyses, and 125 additional species which served for inferring a taxonomically balanced phylogeny and to inform calibrations.

### DNA Amplification, Phylogenetic Reconstruction and Divergence Times Estimation

Out of the 2,612 DNA sequences used for the phylogenetic analyses, 2,490 were retrieved from GenBank, and 122 were produced for this study (Table S1). We included five mitochondrial (*12S*; *16S*; cytochrome b, *cytb*; and NADH dehydrogenases 2 and 4, *ND2* and *ND4*) and six nuclear markers (the acetylcholinergic receptor M4, *ACM4*; the oocyte maturation factor MOS, *C-MOS*; the melanocortin 1 receptor, *MC1R*; neurotrophin-3, *NT3*; and the recombination activating genes 1 and 2, *RAG1* and *RAG2*). To generate the new sequences, genomic DNA was extracted from ethanol-preserved tissue samples using the SpeedTools Tissue DNA Extraction kit (Biotools, Madrid, Spain). Primers and PCR conditions used for the amplification of all genetic fragments are summarized in Table S2. We checked the chromatographs and assembled the contigs in Geneious v.7.1.9 (Kearse et al. 2012). Heterozygous positions in the nuclear genes were coded according to the IUPAC ambiguity codes. We used a supermatrix approach, concatenating all the genetic markers for the 578 samples. The sequence alignments were produced with the online software MAFFT (Katoh et al. 2018) and the concatenated dataset had a length of 12,049 base pairs. We reconstructed the phylogenetic relationships and estimated the divergence times in a single Bayesian inference analysis using the software BEAST2 (Bouckaert et al. 2019). We used 21 calibration points (fossils, biogeographic events, and secondary calibrations), avoiding fixing nodes of potential interest for the biogeographic discussion. Information on calibration points is available in Table S3. Additionally, we constrained higher-level clades to match one of the most recent and supported topologies of squamates published (Streicher and Wiens 2017). We tried different partition schemes, including those resulting from preliminary analyses with PartitionFinder (Lanfear et al. 2016) and ModelFinder (Kalyaanamoorthy et al. 2017) as implemented in IQ-Tree (Nguyen et al. 2015), as well as separating the mitochondrial and the nuclear genes in two different partitions. But all of them failed to converge in the BEAST2 analyses. Thus, we performed five individual runs of 100 million generations each, sampling every 10,000 generations, with a Yule process tree prior, a single partition, and a relaxed lognormal clock model to at least partially account for heterogeneity across different gene fragments. We used the reversible-jump (RB) method of Bouckaert et al. (2013) to sample across multiple substitution models and accommodated the among-site rate heterogeneity using a discretized Gamma distribution with four categories. We used Tracer v.1.7 (Rambaut et al. 2018) to evaluate the stationarity of the chains and check for sufficient effective sample sizes (ESS > 200), as well as convergence between independent runs. We combined the posterior distributions with LogCombiner, discarding 30% of the posterior trees as burn-in. We obtained the maximum clade credibility tree with median heights using TreeAnnotator (both softwares provided within the BEAST2 package).

### Biogeographic Area Delimitation and Ancestral Range Estimation

We used the R package BioGeoBEARS (Matzke 2013) to reconstruct ancestral ranges for each of the 21 monophyletic groups of interest separately, excluding outgroups (see *Taxon sampling and definition of groups of interest*). We assigned each species to their corresponding biogeographic range as a discrete character, delimiting five different areas (Fig. 1): Africa (all the African continent west of the Sinai Peninsula, including Madagascar), Arabia (the Arabian Peninsula south of Jordan and Iraq, including the Socotra Archipelago and Kuwait), Eastern Mediterranean (Lebanon, Syria and Turkey), Asia/Oceania (east of Iraq), and Europe. We defined a ‘contact zone’ in the surroundings of the Arabian Peninsula, consisting of the Sinai Peninsula, Israel, Jordan and Iraq. This allowed us to categorize species mainly distributed in Arabia or Africa and also slightly entering this neutral zone but not the other biogeographic region (i.e., Africa or Arabia, respectively). Species present in more than one of those areas were assigned correspondingly. The maximum number of areas per node was set to two. We performed ancestral reconstructions using the following models in BioGeoBEARS: Dispersal-Extinction-Cladogenesis (DEC; Ree and Smith 2008), DIVALIKE (Ronquist 1997) and BAYAREA (Landis et al. 2013). The *j* parameter representing founder event speciation (Matzke 2014) was excluded from each model due to potential statistical problems (pathological support for models including *j*: Ree and Sanmartín 2018). Fitted models were subsequently selected according to the Akaike information criterion, correcting for small sample size (AICc; Akaike 1973). We extracted ancestral states information at the nodes with the BioGeoBEARS functions *get_ML_probs* and *get_ML_states_from_relprobs*. For each node, we selected the state with the highest marginal probability.

**Figure 1.**
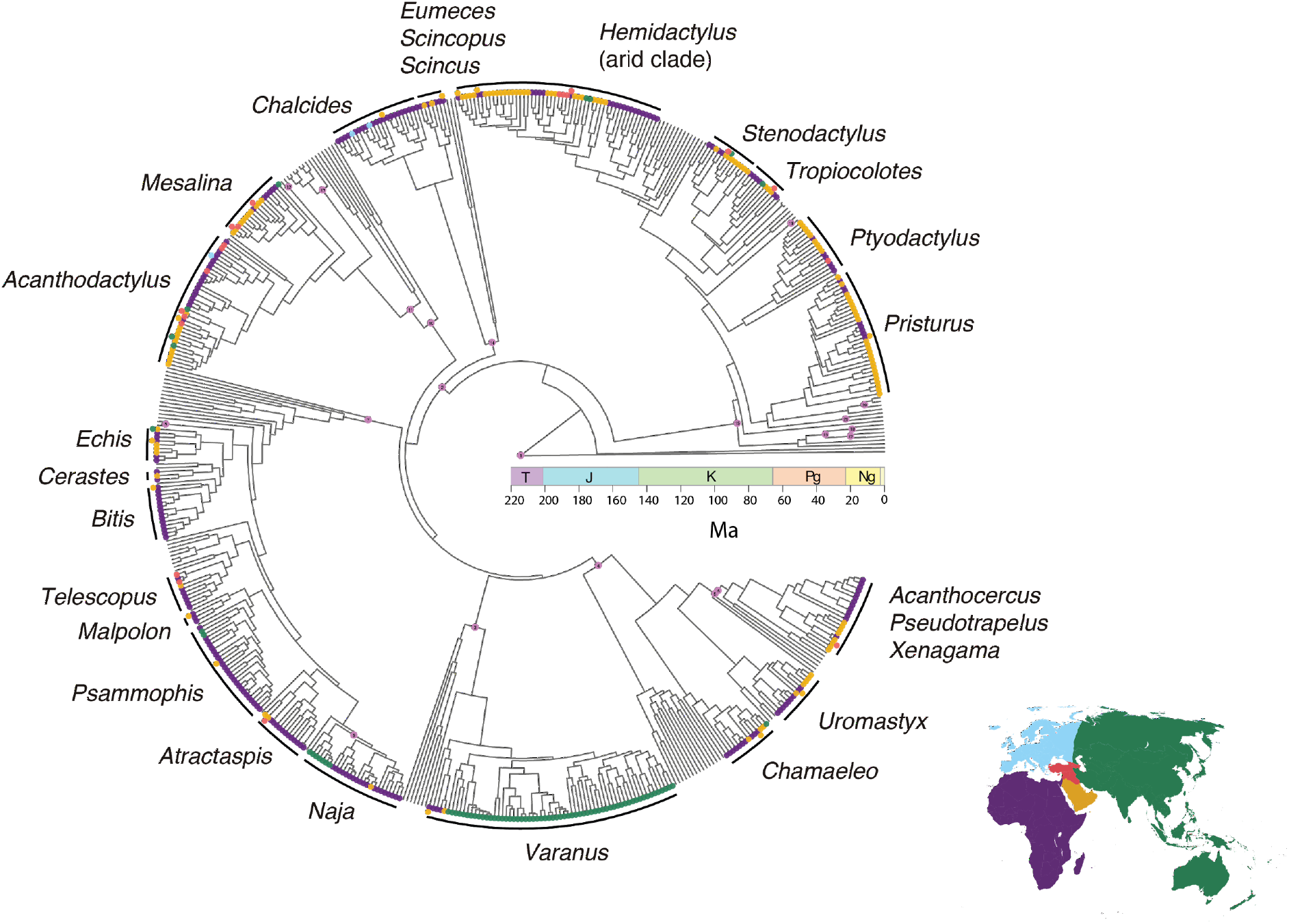
Squamate phylogenetic tree with tips colored by area and calibration points displayed on the corresponding nodes. All the clades used for the biogeographic assessment are indicated. T, Triassic; J, Jurassic; K, Cretaceous; Pg, Paleogene; Ng, Neogene; Ma, millions of years ago. For a more detailed version, including species names and the numbers on calibration points corresponding to Table S3, see Figure S1.

### Biogeographical Assessment and Incidence of Biogeographic Events Through Time

We tracked the counts of the following types of biogeographic events through time: i) dispersal: parent node present in Africa or Arabia, and one of the descendant nodes present in the other area (Arabia or Africa, respectively); ii) vicariance: parent node widespread (Africa+Arabia) and each of the descendant nodes in a different area; and iii) extirpation or range contraction: parent node widespread and one descendant node only present in Africa or Arabia. As pointed out by Klaus et al. (2016), accounting for uncertainty in phylogenetic calibrations of divergence times is essential for the interpretation of biogeographical reconstructions. Therefore, we adopted their approach and included a time range for each biogeographic event, rather than a single time point. We then used the overlap of intervals to estimate the incidence of biogeographic events between Africa and Arabia through time.

For vicariance events, we accounted for temporal uncertainty by including the 95% highest posterior density (HPD) interval of each node inferred to have undergone vicariance between Africa and Arabia. For dispersal and range contraction events, we used time ranges defined as the length of the branch during which the dispersal took place according to our ancestral reconstruction. Combining all clades, we obtained the biogeographic incidence through time by summing the overlapping biogeographic events in each million-year interval. Our metric is analogous to the Maximal number of observed Dispersal Events per million year (MDE) utilized by Klaus et al. (2016) but includes vicariance and extirpation events besides dispersal.

### Model Adequacy and Deviations from the DEC Model

We sought to quantify the extent to which the observed biogeographic history deviates from that expected from the best-fit model and the inferred phylogenetic history alone. That is, we aimed to identify whether (and when) forces beyond those modeled within the DEC framework led to deviations from model expectations. To do so, we simulated the biogeographic history for each of our clades of interest along their respective phylogenies to obtain an expected distribution of biogeographic events through time. We simulated 1,000 per-clade biogeographic histories, running the function *simulate_biogeog_history* in BioGeoBEARS, using the phylogenetic trees, states at the root, empirical transition matrices, and cladogenesis models for each clade of interest. As we did for the observed biogeographic history, we overlapped the temporal uncertainty of biogeographic events in the simulated biogeographic histories and built 1,000 lines of biogeographic incidence through time by summing the number of events in each million-year period. Then we generated a distribution of the expected number of events through time by calculating the 95% confidence interval of the number of simulated biogeographic events per million-year interval. By comparing the number of observed events through time to those expected under the inferred model, we identified specific time periods when the observed number of biogeographic events significantly differed from that expected under the given model and phylogeny alone. We conducted this comparison for all kinds of biogeographic events together and also separately for dispersal in each direction, extirpation from each area (Africa and Arabia), and vicariance. This allowed us to explore and identify time periods during which biogeographic events deviated significantly from model expectations, implying a role of forces external to that modeled by the DEC framework. Figures were produced in the R environment (R Core Team 2019), with the packages ggtree (Yu et al. 2017), treeio (Wang et al. 2020), and the collection of packages within tidyverse (Wickham 2017).

## RESULTS

### Phylogenetic Reconstruction and Divergence Times Estimation

The dated phylogenetic tree of all sampled taxa is shown in Figure 1 (see Figure S1 for details on species names and calibration points). The topology within our clades of interest and crown age estimates do not deviate substantially from previous results (Table S4). Importantly, our approach allows us to include calibration uncertainty in the analyses, a critical step when addressing biogeographic questions (see *Biogeographical assessment* in the Methods section). Further, including all species in the same phylogenetic analysis means that all ages are comparable across clades.

### Ancestral Range Estimation and Biogeographical Assessment

The biogeographic histories for each clade are summarized in Appendix 1, showing the areas reconstructed for each node according to the most supported model (Table S5). Figure 2 shows, for each clade separately, estimated time ranges of biogeographic events involving Africa and Arabia and the type of event (dispersal from Africa to Arabia, dispersal from Arabia to Africa, vicariance, extirpation from Africa, or extirpation from Arabia; see Methods section). Across all clades, the ancestral reconstructions yielded 20 dispersal events from Africa to Arabia and 14 from Arabia to Africa, 26 vicariance events, 8 extirpations from Africa and 6 extirpations from Arabia (Fig. 2). The probability of biogeographic events began to rise between 50 and 40 Ma within some of the oldest clades (e.g. *Acanthodactylus*, *Hemidactylus* or *Tropiocolotes*) and was greatest during the period between 23 and 10 Ma (Fig. 2, Fig. 3a). Results also show a decline in the rate at which biogeographic events occur from 10 Ma to the present.

**Figure 2.**
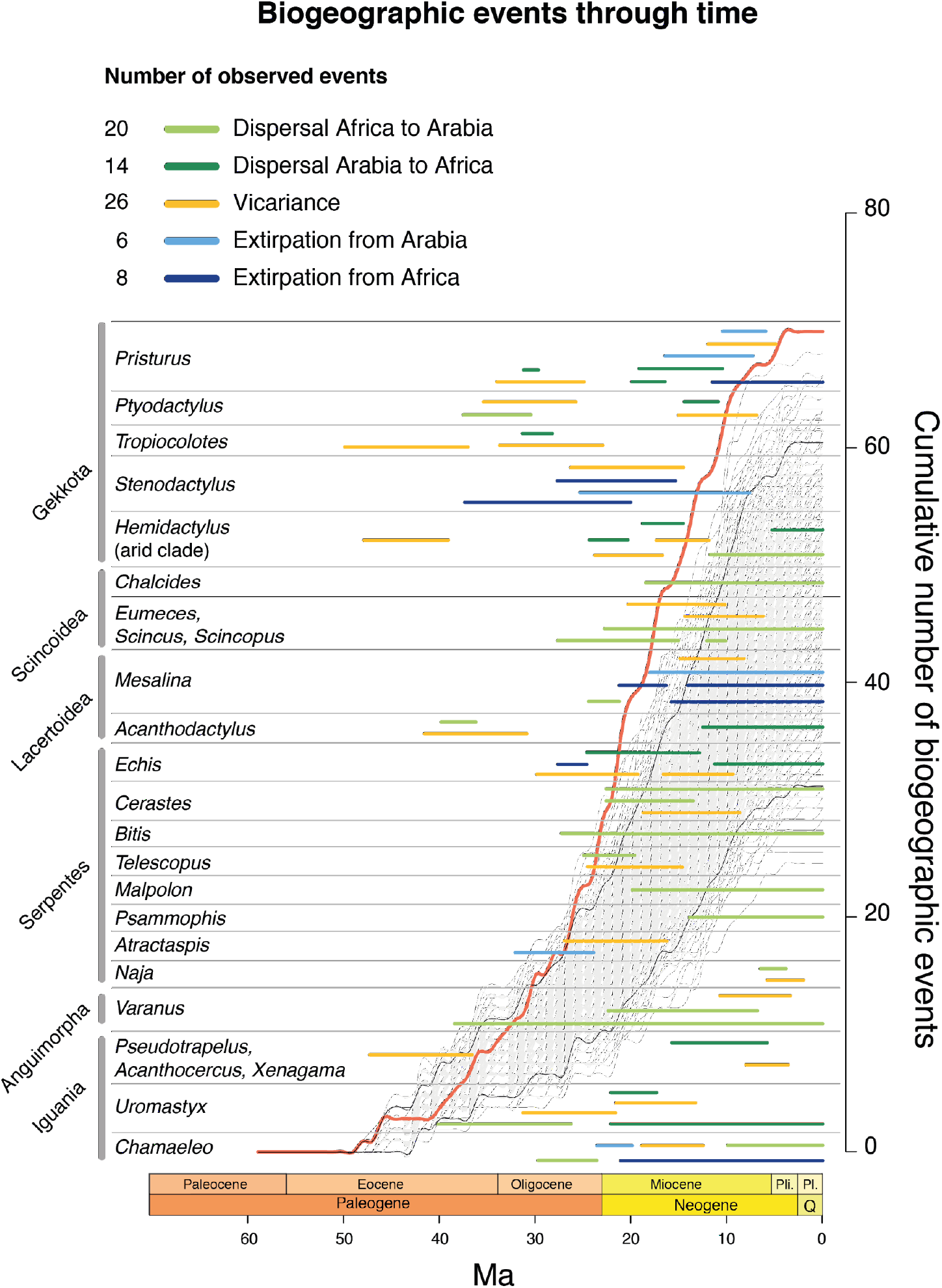
Ages and temporal uncertainty of the biogeographic events involving Africa and Arabia inferred by our ancestral reconstruction for each group of interest. In the background, the red line shows the cumulative observed number of biogeographic events through time, the gray lines show the distribution of the cumulative expected number of events from the simulated biogeographic histories, and the black lines enclose the 95% confidence interval of that distribution. Pli., Pliocene; Pl., Pleistocene; Q, Quaternary.

**Figure 3.**
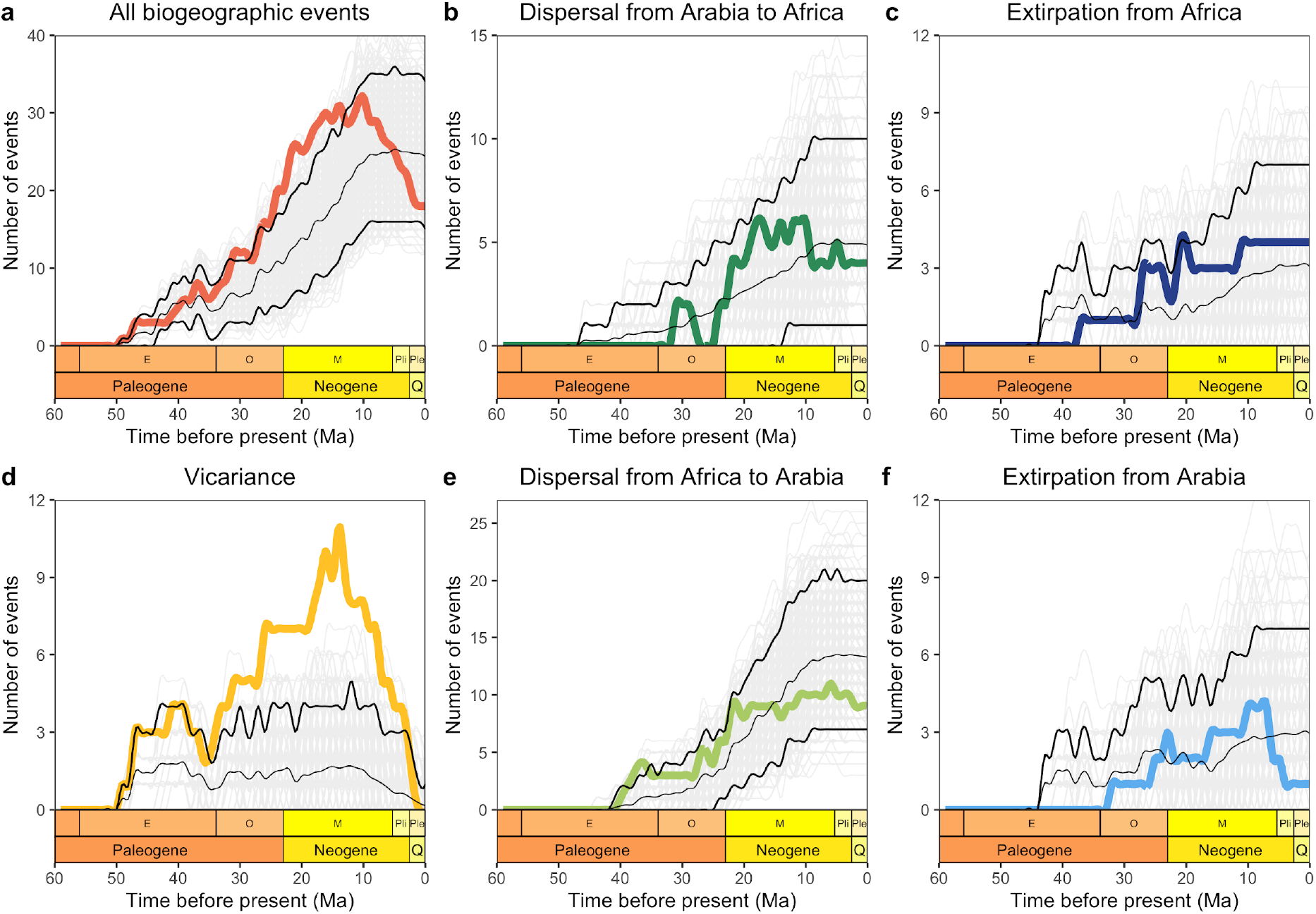
Observed (colored lines) and expected (background gray lines) incidence of biogeographic events through time. Thick black lines delimit the 95% interval of events simulated, while the thin black line represents the average. a) total number of events; b-f) different types of events separated. Colors correspond to Figure 2. Deviations of the observed number of events from the expected distribution envelope indicate the inadequacy of the DEC model. Most notably, the peak in vicariance events during the Oligocene and Miocene suggests forces external to those modeled in the DEC framework have played a role in shaping squamate biogeographic patterns in Afro-Arabia.

### Incidence of Biogeographic Events Through Time and Model Adequacy

By summing the number of overlapping biogeographic events across all clades in each million-year interval, we can quantify how the frequency of events in the biogeographic history has changed through time. However, the dynamics of species accumulation alone can lead to complex patterns in the timing of these events, as seen when we reconstruct events using the 1,000 simulated biogeographic histories. We can then compare the actual reconstructed events to this expected distribution of biogeographic events through time, considering only the shape of the phylogeny, the number of lineages through time, and the best-fit biogeographic model. This is shown in Figure 3, including all types of events together (Fig. 3a) and each type separately (Fig. 3b-f). Any deviation of the observed number of events from the expected distribution indicates the inadequacy of the DEC model to fully capture the temporal dynamics of biogeographic events.

When summing across all types of biogeographic events, we find that the observed number of events does not differ substantially from the model expectation, with the exception of the period between 23 and 15 Ma, during which we observed slightly more incidence than expected (Fig. 3a). In contrast, looking at each type of biogeographic event separately gives a much more nuanced understanding. Most notably, the incidence of vicariance events was considerably higher than expected beginning around 30 Ma until about 3 Ma (Fig. 3d), a likely explanation for the excess of biogeographic events observed around the same time (Fig. 2).

The number of observed events of dispersal (both from Arabia to Africa and from Africa to Arabia; Figs. 3b and 3e, respectively) and range contraction (Figs. 3c and 3f) have not been significantly different than expected. The levels of dispersal in both directions suggest a potential asymmetry of dispersal rates and exchange among the two regions. Indeed, the interchange between Africa and Arabia has not been symmetrical (Fig. 4). Dispersal started earlier from Africa to Arabia (until around 32 Ma all dispersal happened in that direction). Later, through the Oligocene and the Early Miocene, the relative rate of dispersal from Arabia to Africa intensified, although it never matched the levels of dispersal from Africa to Arabia, which have always been predominant. This asymmetry, however, is indistinguishable from that expected given the phylogeny and fitted biogeographic model.

**Figure 4.**
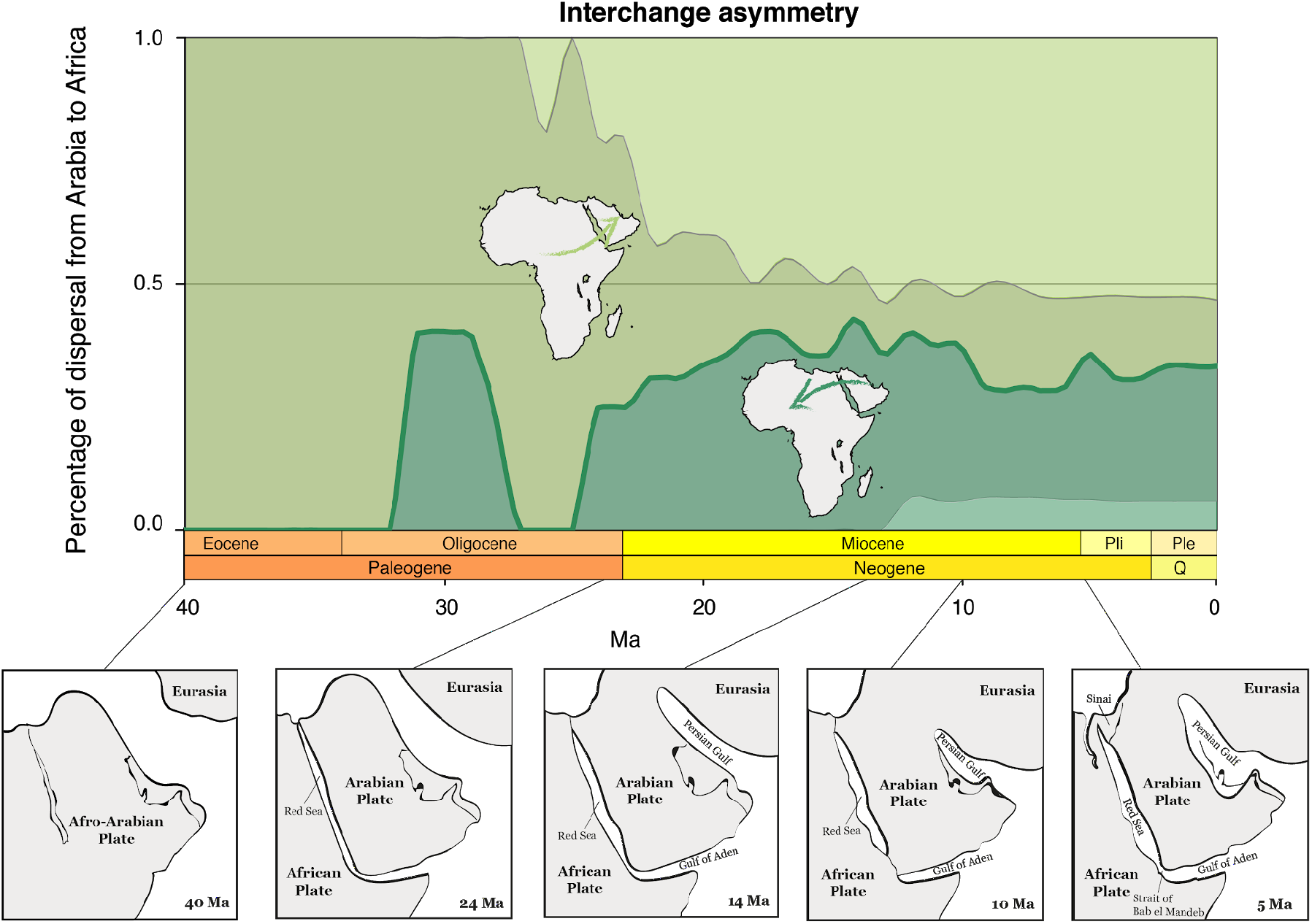
Interchange asymmetry between Africa and Arabia over time. Green shaded areas represent, in percentage, the relative intensity of observed dispersal from Africa to Arabia (up, in light green) and from Arabia to Africa (bottom, in dark green). The dark green line shows the observed proportion of dispersal from Arabia to Africa. The darker shaded area in the background encompasses the 95% interval of the expected distribution of dispersal proportion from Arabia to Africa over time according to the 1,000 biogeographic simulations. Observed asymmetry meets expectations where the green line falls within the 95% of the expected asymmetry (i.e., throughout all the time period shown). Paleogeographic maps adapted from Bosworth et al. 2005.

## Discussion

Over the last few decades, historical biogeography as a field has experienced major theoretical and methodological advances, resulting in an increasing number of studies that discuss distributional patterns in the context of past environmental processes according to ancestral range estimations. Our novel approach goes one step further and allows us to identify particular periods in the past when biogeographic events happened more or less frequently than predicted by model expectations. That is, we exploit model inadequacies to learn more than by taking fitted models at face value alone. This approach enhances our ability to ask questions central to the study of biogeography, such as what are the historical causes of exceptional historical biogeographic patterns observed in the data and how have those patterns unfolded into current species geographic distributions?

Our results show a substantial contrast between the number of reptile biogeographic events involving Africa and Arabia observed in the last 40 million years and the expected number of such events simulated under the best-fit models for the same period. The excess in the incidence of vicariance events between Africa and Arabia during the Oligocene and the Miocene (approximately from 30 to 5 Ma; Fig. 3d) is of particular interest. We hypothesize this pattern is driven by the movement of the Arabian and the African plates (Fig. 4). These two plates, once connected in a single landmass, started splitting around 40-30 Ma (Bosworth et al. 2005; Scotese 2014*b*), initially with the uplifting of mountain chains along the plates’ limits and eventually resulting in the establishment of the Red Sea around 25 Ma (Ghebreab 1998). This fragmentation is considered to be a major driver of the biogeographic history of taxa currently present in the Afro-Arabian area (Pook et al. 2009; Metallinou et al. 2012; Šmíd et al. 2013; Portik and Papenfuss 2015; Tamar et al. 2016*a*, 2018). Further, the aridification of the region, that began around the middle Miocene (Flower and Kennett 1994), has been suggested to have a significant role in promoting vicariant speciation (Metallinou et al. 2015). Our results suggest this interval of tectonic and climatic instability had a crucial role in shaping the observed biogeographic patterns of Afro-Arabian reptiles, with many clades undergoing vicariance as well as dispersing out of their place of origin from the late Eocene through the middle Miocene (Fig. 2). Consequently, the observed increase in vicariance events beyond DEC model expectations from 30 to 10 Ma (Fig. 3d) is likely related to these geologic and environmental processes involving Africa and Arabia, which is also widely supported in the literature (Šmíd et al. 2013; Tamar et al. 2016*a*).

The fact that range contraction, and especially dispersal patterns, do not differ from biogeographic expectations (Fig. 3b, c, d, f) is also worth discussing, given the complex paleogeographic and climatic history of the region. The fragmentation of the Afro-Arabian plate, which culminated at the end of the Oligocene, would lead us to hypothesize a reduction in the observed rate of dispersal from that time onwards. Likewise, the climatic fluctuations that could have imposed environmental barriers between Africa and Arabia since the Miocene (Tchernov 1992; Pickford and Morales 1994) precluding dispersal and promoting range contraction on the regional fauna, would make us expect a reduced incidence of these biogeographic events compared to the simulated history. These results suggest that dispersal and extirpation patterns might have been shaped by the interplay of diversification dynamics and external forces. Namely, the progressive aridification during the Miocene and the Plio-Pleistocene climatic oscillations have been suggested to trigger diversification in Afro-Arabian faunas (Fu 2000; Douady et al. 2003; Guillaumet et al. 2008; Brito et al. 2014; Tamar et al. 2016*a*). This, together with the vicariant speciation promoted by the Afro-Arabian plate fragmentation, may have driven the observed dispersal and extirpation patterns. Nevertheless, these results may also reveal the importance of plate tectonics in the region following the formation of the Red Sea. While the expansion of the Red Sea was acting as a potential vicariant agent, the landmasses came into contact several times through different time periods. The connection of Eurasia and Afro-Arabia through the *Gomphotherium* landbridge around 18 Ma and the later collision of the Arabian plate with Eurasia around 15 Ma (Rögl 1999; Jolivet and Faccenna 2000; Bosworth et al. 2005) have played an important role as a potential dispersal route in the Miocene faunal interchange (Barry et al. 1991; Joger 1991; Nakaya 1994; Kappelman et al. 2003; Harzhauser et al. 2007; Tamar et al. 2016*b*).

This dispersal route has been permanent since the middle Miocene, connecting Africa and Arabia through the Sinai Peninsula. Likewise, the global decline in sea level during the late Miocene (Flower and Kennett 1994) resulted in a temporary southwestern landbridge between the Arabian Peninsula and the Horn of Africa that was in place from approximately 10 to 5.3 Ma (Bosworth et al. 2005). It has been suggested that this land connection might have served as a dispersal route from Africa to Arabia and vice versa (Tassy 1999; Portik and Papenfuss 2012; Šmíd et al. 2013) until the reopening of the Bab el Mandeb Strait during the Pliocene, which has persisted until the present. Furthermore, aridification trends may have contributed to the dispersal of reptile taxa (Tamar et al. 2018). Here we found evidence that the landbridge that emerged as a consequence of the connection between the African, Arabian and Eurasian plates served as a permanent dispersal corridor through the entire geologic history after the Red Sea opening, suggesting an environmental homogeneity between the Sahara and the Arabian deserts. This is consistent with the lack of difference between observed and expected patterns, both for dispersal and range contraction events. Another interesting aspect of our results is that the reptile interchange between Africa and Arabia shows a clear asymmetry (Fig. 4). The predominant direction of the interchange, from Africa to Arabia, suggests a scenario of classic island biogeography source-sink dynamics, where Africa has primarily acted as the ‘mainland’ source of colonizing species into Arabia. After the establishment of the Red Sea at the end of the Oligocene, the relative levels of dispersal in both directions have remained stable, which further highlights the role of the dispersal corridor in shaping biogeographic patterns and the environmental homogeneity on both sides of the Red Sea.

Overall, our study corroborates the importance of the unstable geologic and climatic history of Afro-Arabia as a key factor to explain the evolutionary history of the fauna in this region. Our results suggest distinct roles of tectonic and climatic processes in different periods since the Oligocene. The fragmentation of the Afro-Arabian plate seems to be the main driver of biogeographic patterns through the Oligocene and Miocene. Additionally, the landmass movement after the fragmentation, together with diversification dynamics driven by climatic fluctuations, appear to be more relevant in explaining the biogeographic patterns from the middle Miocene onwards. In future studies, integrating data from other taxa and from the fossil record may allow us to study these patterns in a more comprehensive way, and search for shared evolutionary patterns among groups across the tree of life.

The use of null models and adequacy testing is common in areas of evolutionary biology and ecology, such as the study of diversification dynamics (FitzJohn 2012; Beaulieu and O’Meara 2016; Caetano et al. 2018). Our approach, complementary to recent advances in the analysis of community assembly (Hua and Bromham 2020), represents a growing recognition of the potential inadequacy of models and the need to test for it. Here we show that model inadequacy can help us to better understand complex biological patterns. By quantifying model adequacy through time, we have identified specific time periods during which unmodeled processes (i.e. tectonic plate movement) may have led to unexpectedly high rates of vicariance. Our approach made it possible to disentangle extrinsic and intrinsic factors that may have determined the evolutionary history of the groups. Ultimately, such a perspective can provide an important insight from the past to help us face the challenges imposed by the intense changes in our planet’s present and future environmental conditions.

## DATA AND CODE AVAILABILITY

Phylogenetic tree, distributional data, and R code for the analyses and figures are available on GitHub: https://github.com/Cicuendez/Afro-Arabian_biogeo

## ACKNOWLEDGEMENTS

We are very grateful to M. Metallinou, K. Tamar, M. Simó-Riudalbas, L. Machado, J.G. Porta, D. Gonçalves, J.C. Brito, J. Harris, P-A. Crochet, Ph. Geniez, T. Mazuch, J. Moravec, L. Kratochvíl, and B. Burriel for their help in different aspects of the work. We also wish to thank Iris Menéndez for her insightful comments on the text and figures.

## FUNDING

This study was funded by grants CGL2015-70390-P and PGC2018-098290-B-I00 from the Ministerio de Economía y Competitividad, Spain (cofunded by FEDER) and grant 2017-SGR-00991 from the Secretaria d’Universitats i Recerca del Departament d’Economia i Coneixement de la Generalitat de Catalunya to SC. HTC was funded by an FPI grant from the Ministerio de Economía y Competitividad, Spain (BES-2016-078341). JŠ was supported by the Czech Science Foundation (GACR, project number 18–15286Y) and the Ministry of Culture of the Czech Republic (DKRVO 2019–2023/6.VII.c, 00023272).

## Notes

https://github.com/Cicuendez/Afro-Arabian_biogeo

### Competing Interest Statement

The authors have declared no competing interest.

