## Supplementary figures and images for "Reconstructing Squamate Biogeography in Afro-Arabia Reveals the Influence of a Complex and Dynamic Geologic Past"

### Supplemental Figure 1

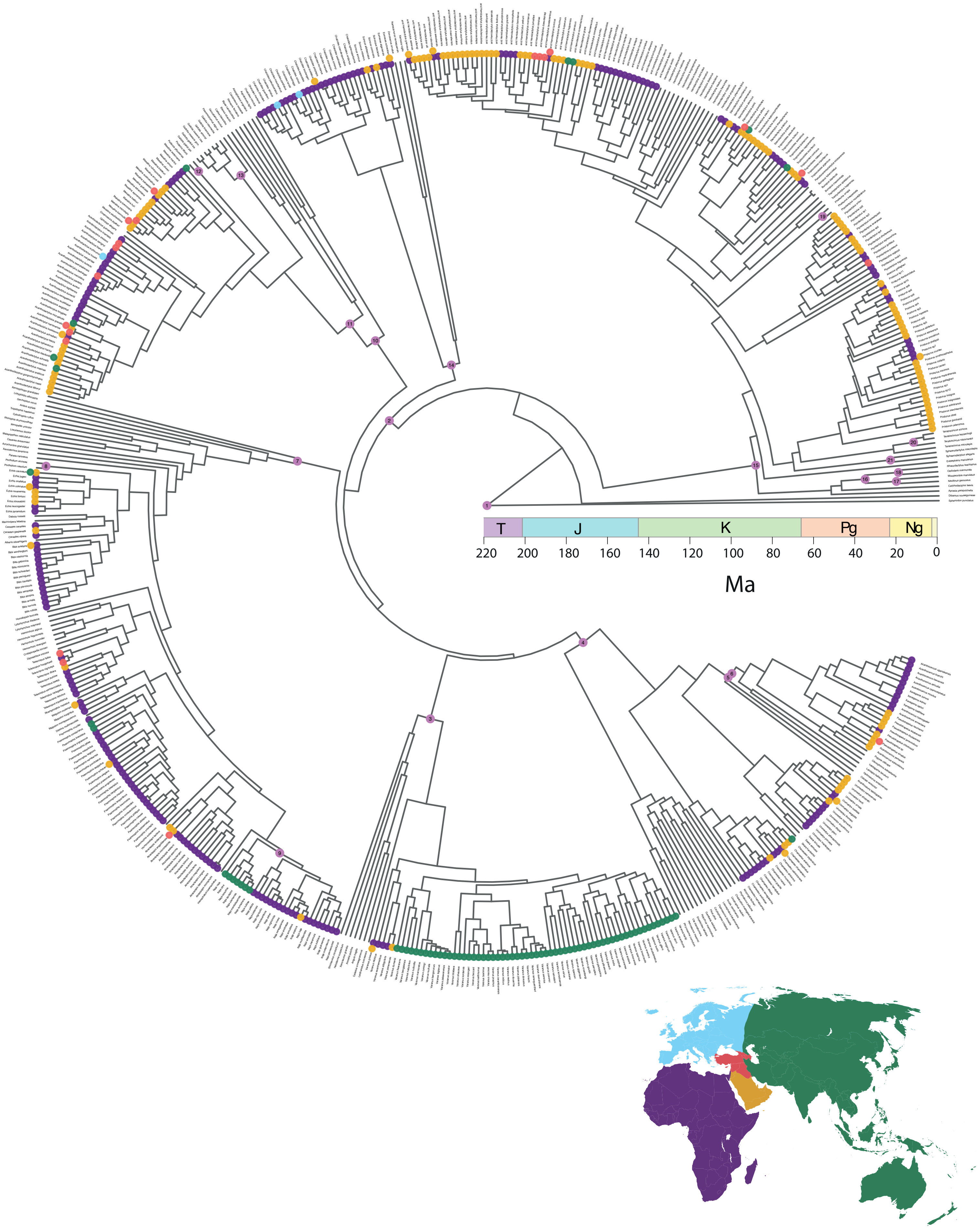
