## Appendix 1 for "Reconstructing Squamate Biogeography in Afro-Arabia Reveals the Influence of a Complex and Dynamic Geologic Past"

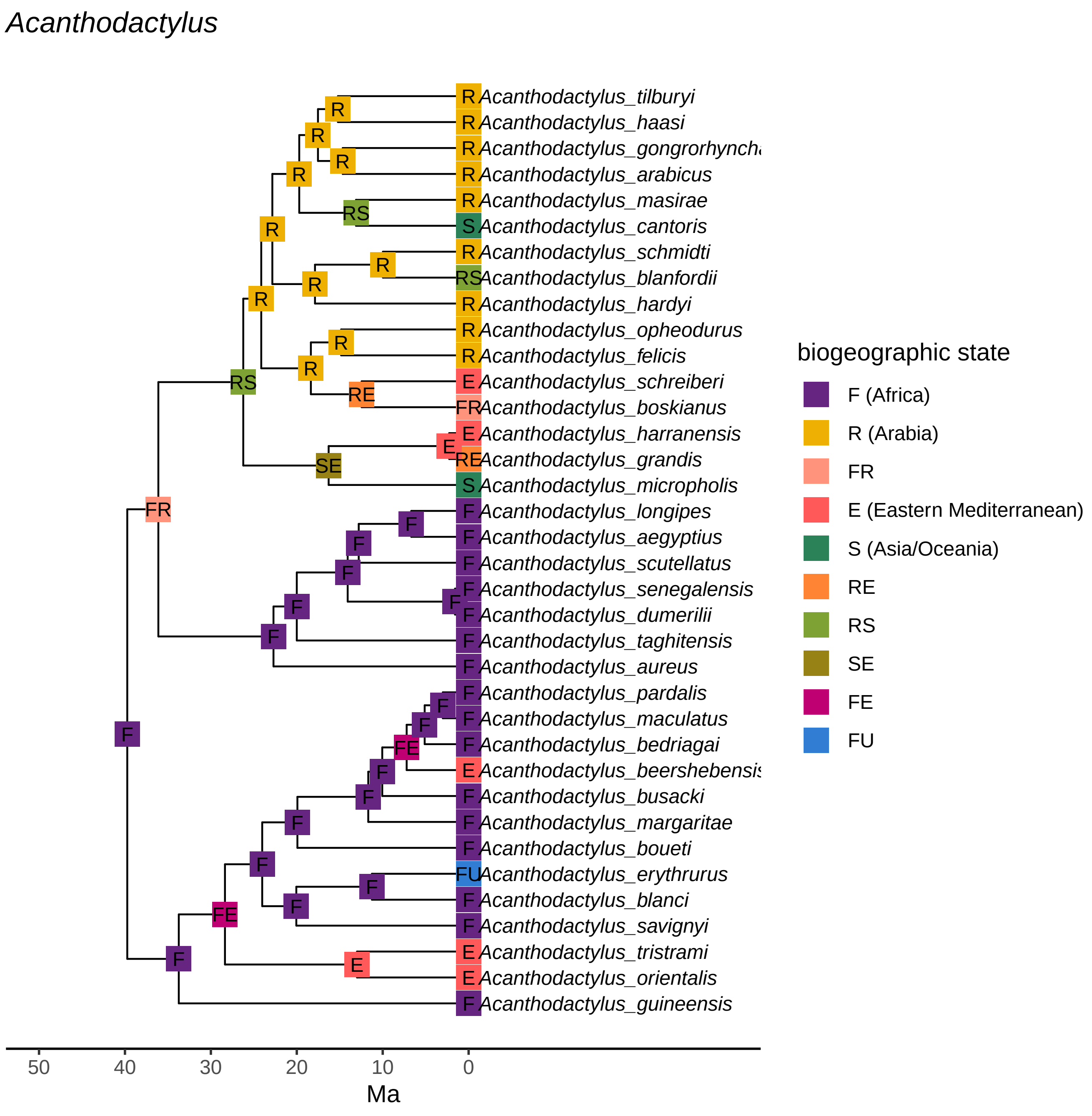
Acanthodactylus
R Acanthodactylus_tilburyi
R Acanthodactylus_haasi
R Acanthodactylus_gongrorhynch
R Acanthodactylus_arabicus
R Acanthodactylus_masirae
S Acanthodactylus_cantoris
R Acanthodactylus_schmidti
RS Acanthodactylus_blanfordii
R Acanthodactylus_hardyi
R Acanthodactylus_opheodurus
R Acanthodactylus_felicis
E Acanthodactylus_schreiberi
FR Acanthodactylus_boskianus
E Acanthodactylus_harranensis
RE Acanthodactylus_grandis
S Acanthodactylus_micropholis
F Acanthodactylus_longipes
F Acanthodactylus_aegyptius
F Acanthodactylus_scutellatus
F Acanthodactylus_senegalensis
F Acanthodactylus_dumerilii
F Acanthodactylus_taghitensis
F Acanthodactylus_aureus
F Acanthodactylus_pardalis
F Acanthodactylus_maculatus
F Acanthodactylus_bedriagai
E Acanthodactylus_beershebensis
F Acanthodactylus_busacki
F Acanthodactylus_margaritae
F Acanthodactylus_boueti
FU Acanthodactylus_erythrurus
F Acanthodactylus_blanci
F Acanthodactylus_savignyi
E Acanthodactylus_tristrami
E Acanthodactylus_orientalis
F Acanthodactylus_guineensis
50
40
30
20
10
0
Ma
biogeographic state
F (Africa)
R (Arabia)
FR
E (Eastern Mediterranean)
S (Asia/Oceania)
RE
RS
SE
FE
FU

*Atractaspis*

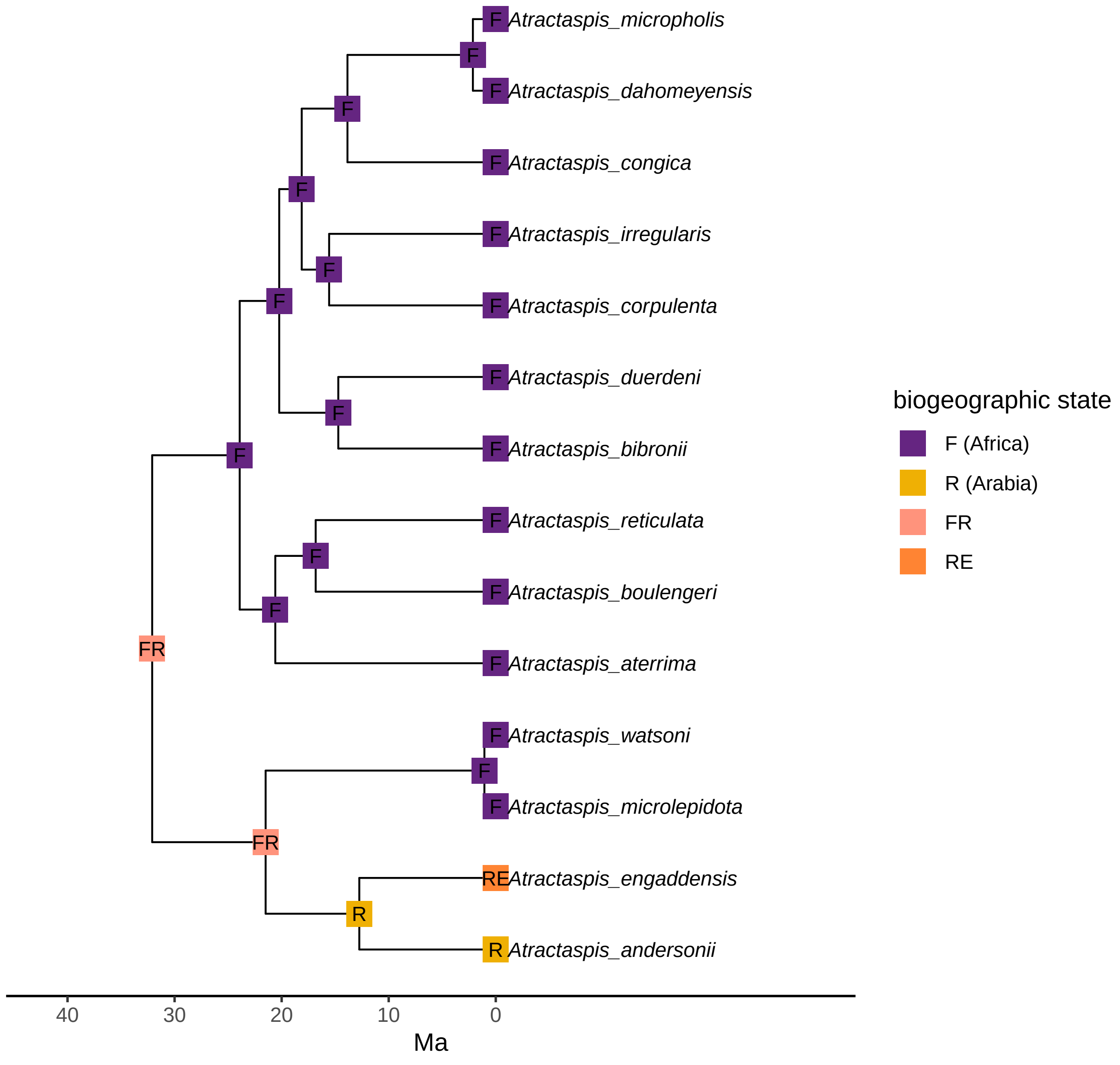

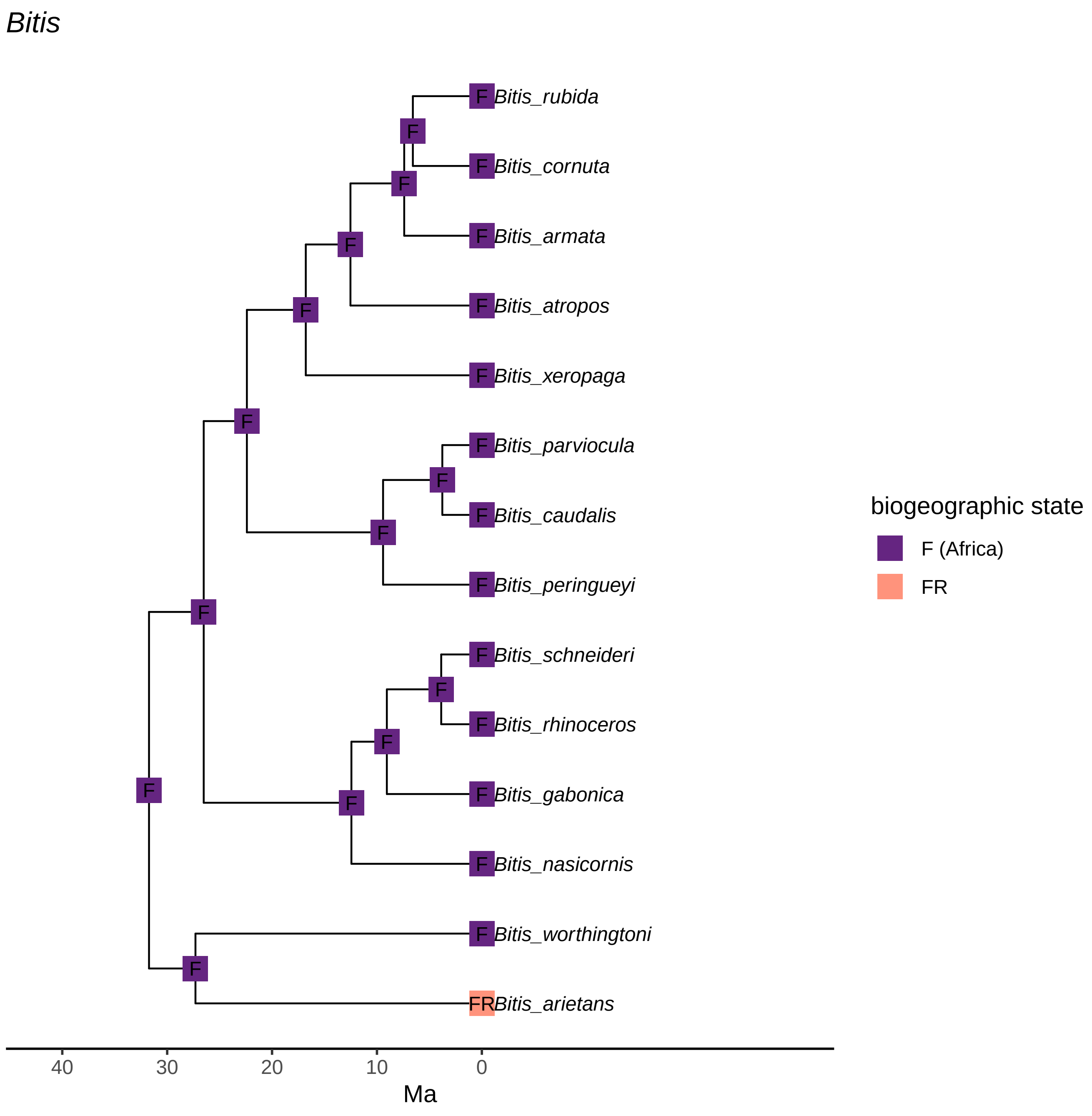

Bitis
Bitis_rubida
Bitis_cornuta
Bitis_armata
Bitis_atropos
Bitis_xeropaga
Bitis_parviocula
Bitis_caudalis
Bitis_peringueyi
Bitis_schneideri
Bitis_rhinoceros
Bitis_gabonica
Bitis_nasicornis
Bitis_worthingtoni
Bitis_arietans
40
30
20
10
0
Ma
biogeographic state
F (Africa)
FR

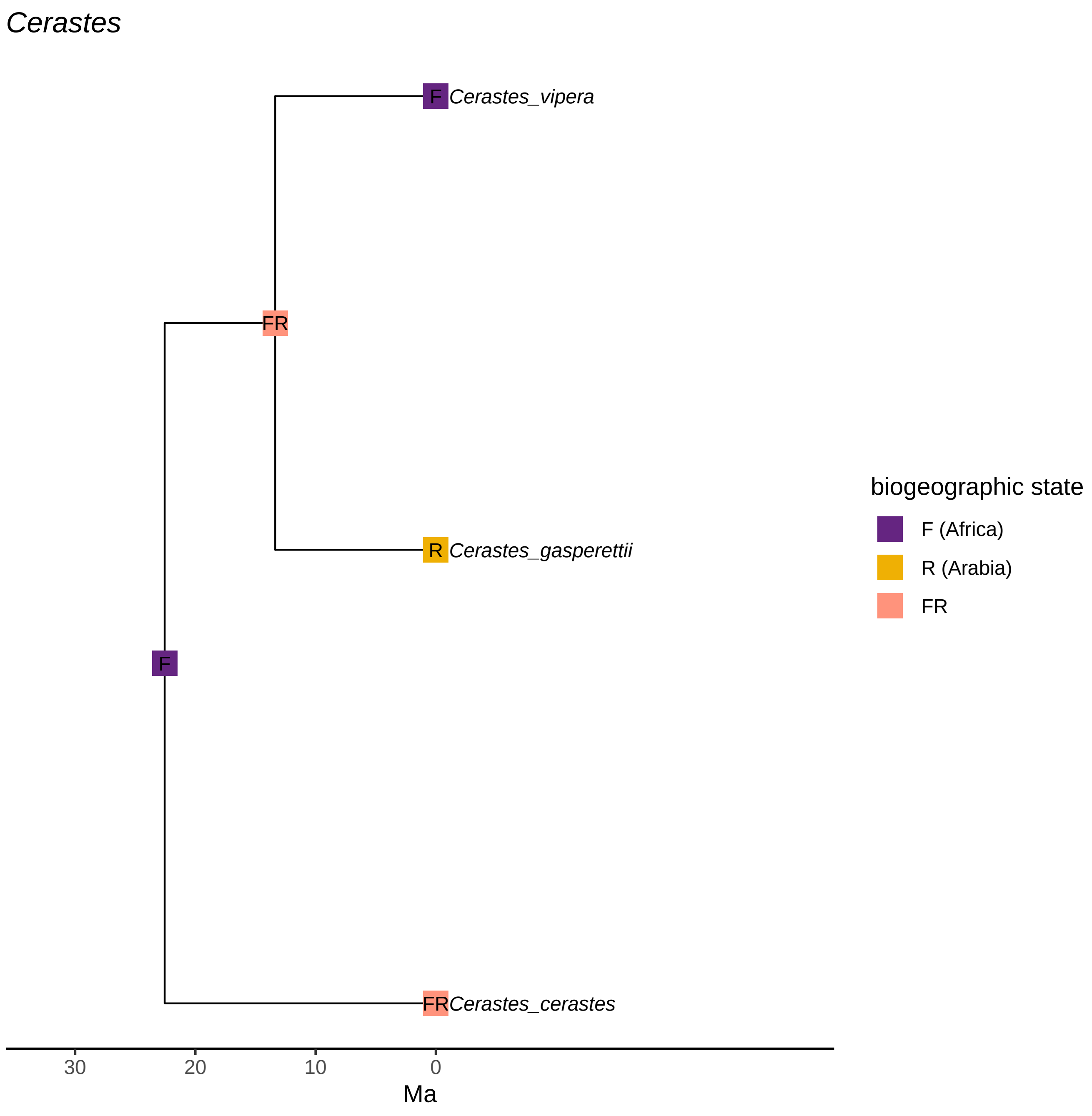
Cerastes
F
Cerastes_vipera
FR
R
Cerastes_gasperettii
F
FR
Cerastes_cerastes
30
20
10
0
Ma
biogeographic state
F (Africa)
R (Arabia)
FR

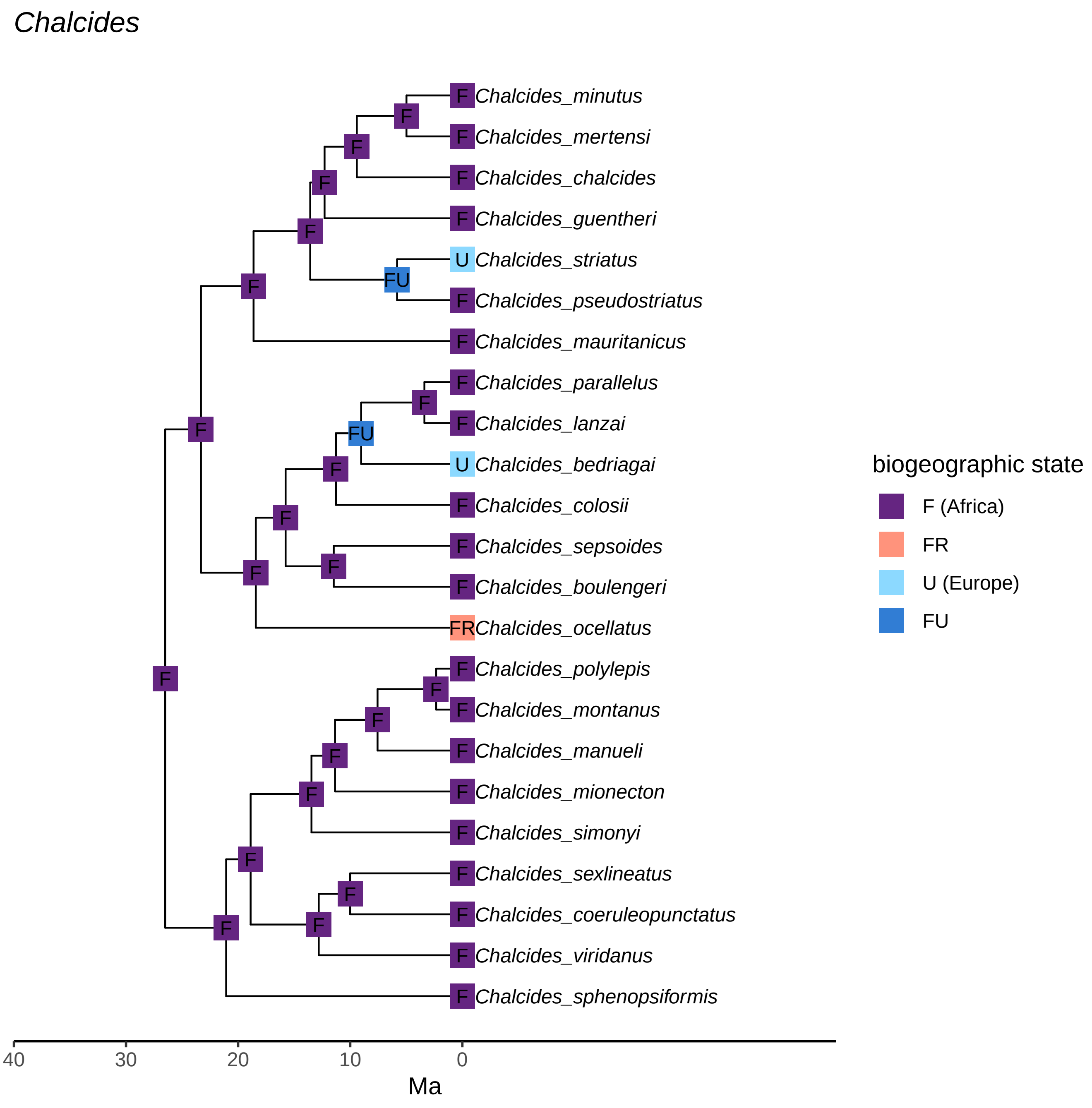
Chalcides
Chalcides_minutus
Chalcides_mertensi
Chalcides_chalcides
Chalcides_guentheri
Chalcides_striatus
Chalcides_pseudostriatus
Chalcides_mauritanicus
Chalcides_parallelus
Chalcides_lanzai
Chalcides_bedriagai
Chalcides_colosii
Chalcides_sepsoides
Chalcides_boulengeri
Chalcides_ocellatus
Chalcides_polylepis
Chalcides_montanus
Chalcides_manueli
Chalcides_mionecton
Chalcides_simonyi
Chalcides_sexlineatus
Chalcides_coeruleopunctatus
Chalcides_viridanus
Chalcides_sphenopsiformis
biogeographic state
F (Africa)
FR
U (Europe)
FU
40
30
20
10
0
Ma

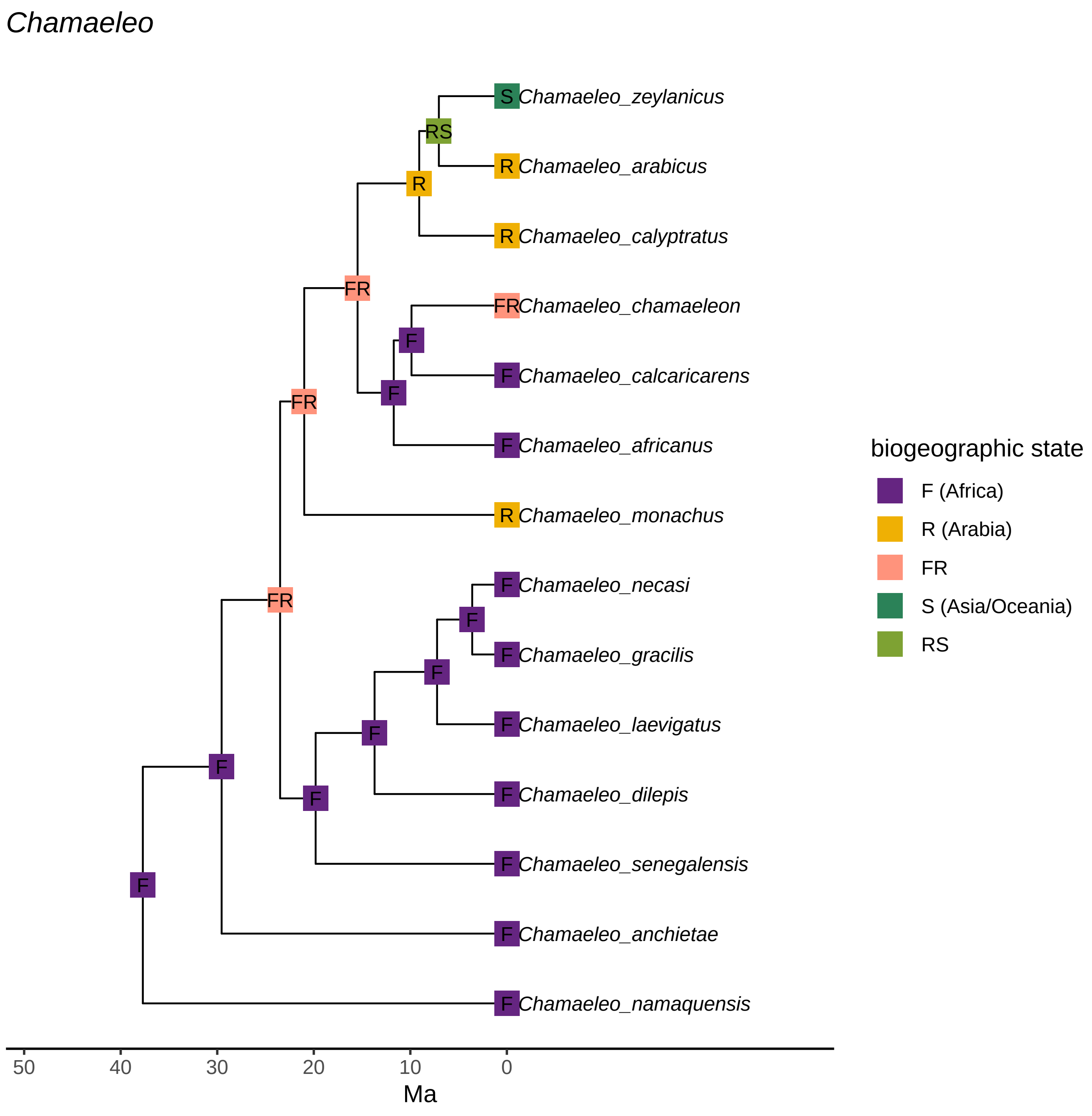
Chamaeleo
S Chamaeleo_zeylanicus
RS
R Chamaeleo_arabicus
R
R Chamaeleo_calyptratus
FR
FR Chamaeleo_chamaeleon
F
F Chamaeleo_calcaricarens
F
FR
F Chamaeleo_africanus
R Chamaeleo_monachus
FR
F Chamaeleo_necasi
F
F Chamaeleo_gracilis
F
F Chamaeleo_laevigatus
F
F
F Chamaeleo_dilepis
F
F Chamaeleo_senegalensis
F Chamaeleo_anchietae
F Chamaeleo_namaquensis
50
40
30
20
10
0
Ma
biogeographic state
F (Africa)
R (Arabia)
FR
S (Asia/Oceania)
RS

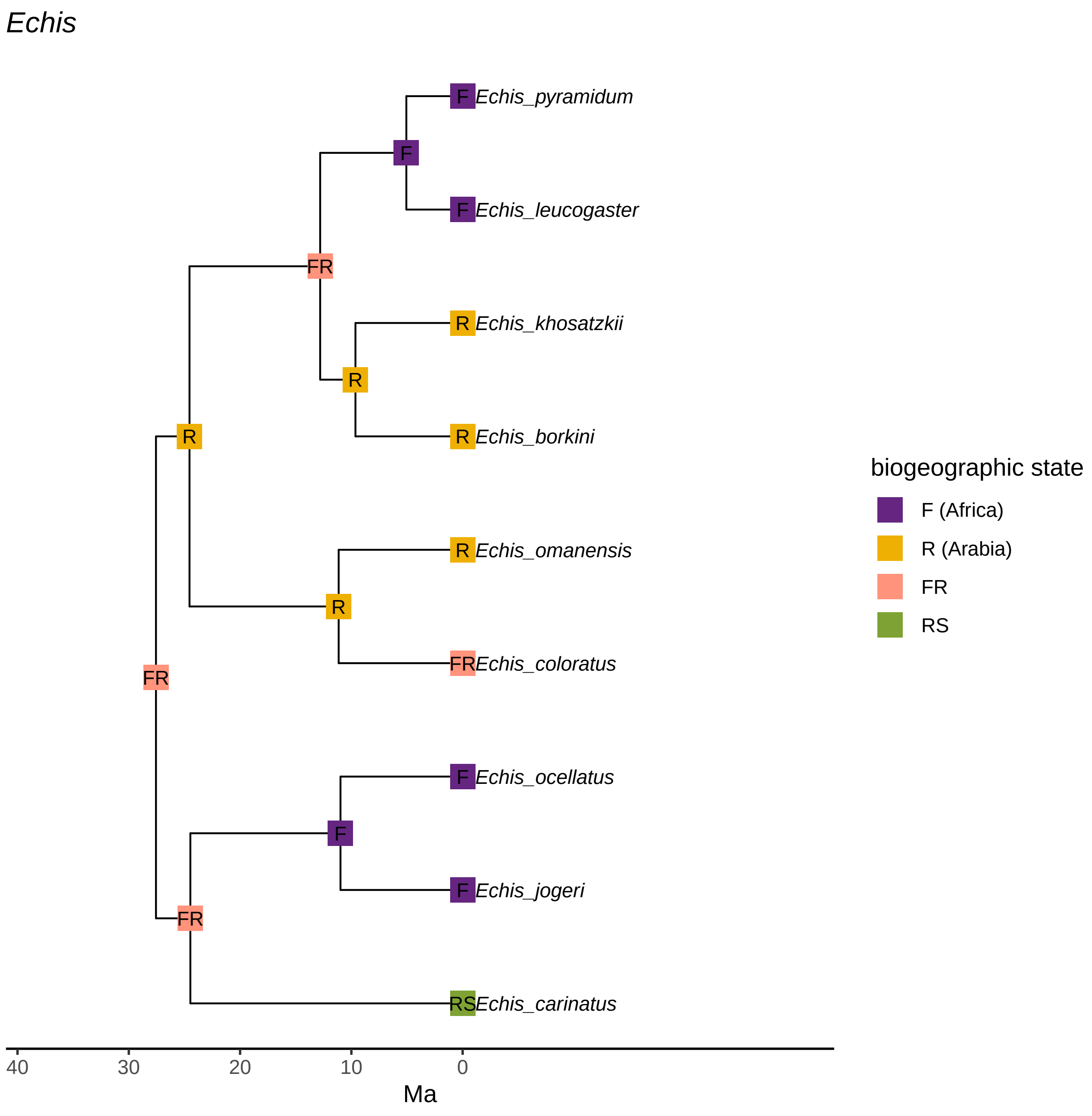
Echis
F
Echis_pyramidum
F
F
Echis_leucogaster
FR
R
Echis_khosatzkii
R
R
R
Echis_borkini
R
Echis_omanensis
R
FR
Echis_coloratus
FR
F
Echis_ocellatus
F
F
Echis_jogeri
FR
RS
Echis_carinatus
40
30
20
10
0
Ma
biogeographic state
F (Africa)
R (Arabia)
FR
RS

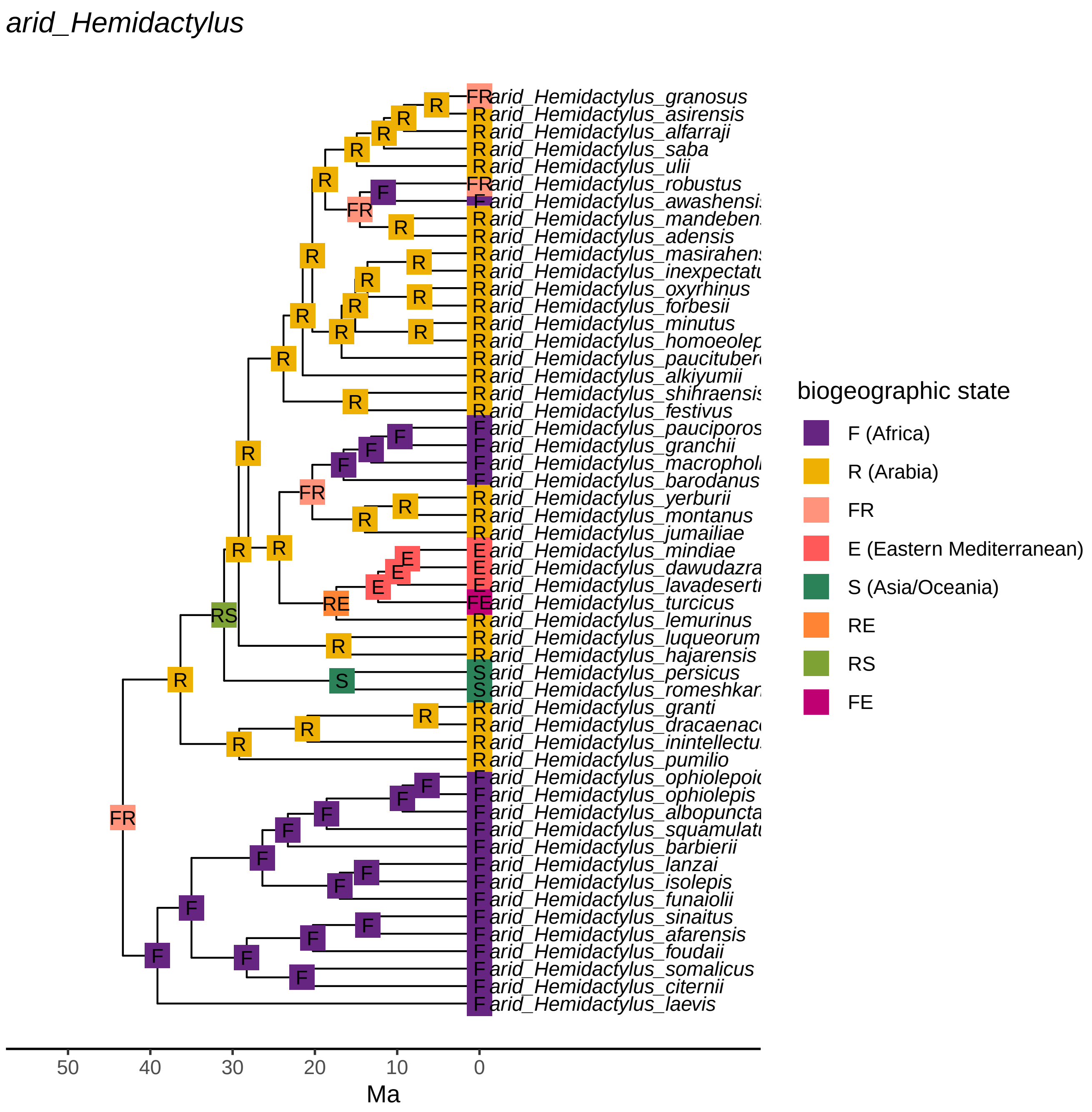
arid_Hemidactylus
arid_Hemidactylus_granosus
arid_Hemidactylus_asirensis
arid_Hemidactylus_alfarraji
arid_Hemidactylus_saba
arid_Hemidactylus_ulii
arid_Hemidactylus_robustus
arid_Hemidactylus_awashensi
arid_Hemidactylus_mandeben
arid_Hemidactylus_adensis
arid_Hemidactylus_masirahen
arid_Hemidactylus_inexpectatu
arid_Hemidactylus_oxyrhinus
arid_Hemidactylus_forbesii
arid_Hemidactylus_minutus
arid_Hemidactylus_homoeolep
arid_Hemidactylus_paucituber
arid_Hemidactylus_alkiyumii
arid_Hemidactylus_shihraensis
arid_Hemidactylus_festivus
arid_Hemidactylus_pauciporos
arid_Hemidactylus_granchii
arid_Hemidactylus_macrophol
arid_Hemidactylus_barodanus
arid_Hemidactylus_yerburii
arid_Hemidactylus_montanus
arid_Hemidactylus_jumailiae
arid_Hemidactylus_mindiae
arid_Hemidactylus_dawudazra
arid_Hemidactylus_lavadeserti
arid_Hemidactylus_turcicus
arid_Hemidactylus_lemurinus
arid_Hemidactylus_luqueorum
arid_Hemidactylus_hajarensis
arid_Hemidactylus_persicus
arid_Hemidactylus_romeshkan
arid_Hemidactylus_granti
arid_Hemidactylus_dracaenac
arid_Hemidactylus_inintellectu
arid_Hemidactylus_pumilio
arid_Hemidactylus_ophiolepoi
arid_Hemidactylus_ophiolepis
arid_Hemidactylus_albopuncta
arid_Hemidactylus_squamulatu
arid_Hemidactylus_barbierii
arid_Hemidactylus_lanzai
arid_Hemidactylus_isolepis
arid_Hemidactylus_funaiolii
arid_Hemidactylus_sinaitus
arid_Hemidactylus_afarensis
arid_Hemidactylus_foudaii
arid_Hemidactylus_somalicus
arid_Hemidactylus_citernii
arid_Hemidactylus_laevis
50 40 30 20 10 0
Ma
biogeographic state
F (Africa)
R (Arabia)
FR
E (Eastern Mediterranean)
S (Asia/Oceania)
RE
RS
FE

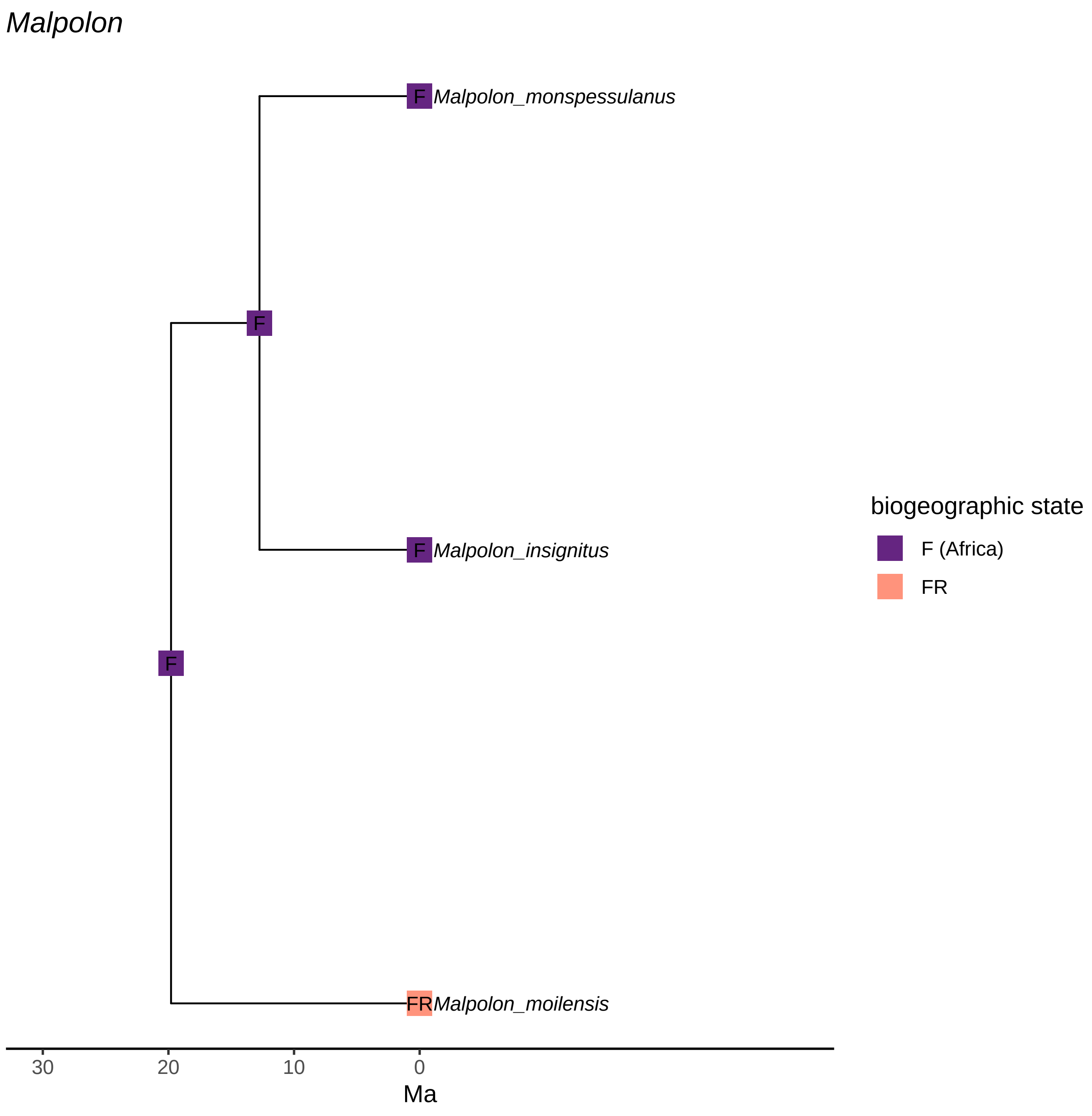
Malpolon
F
Malpolon_monspessulanus
F
F
Malpolon_insignitus
F
FR
Malpolon_moilensis
30
20
10
0
Ma
biogeographic state
F (Africa)
FR

*Mesalina*

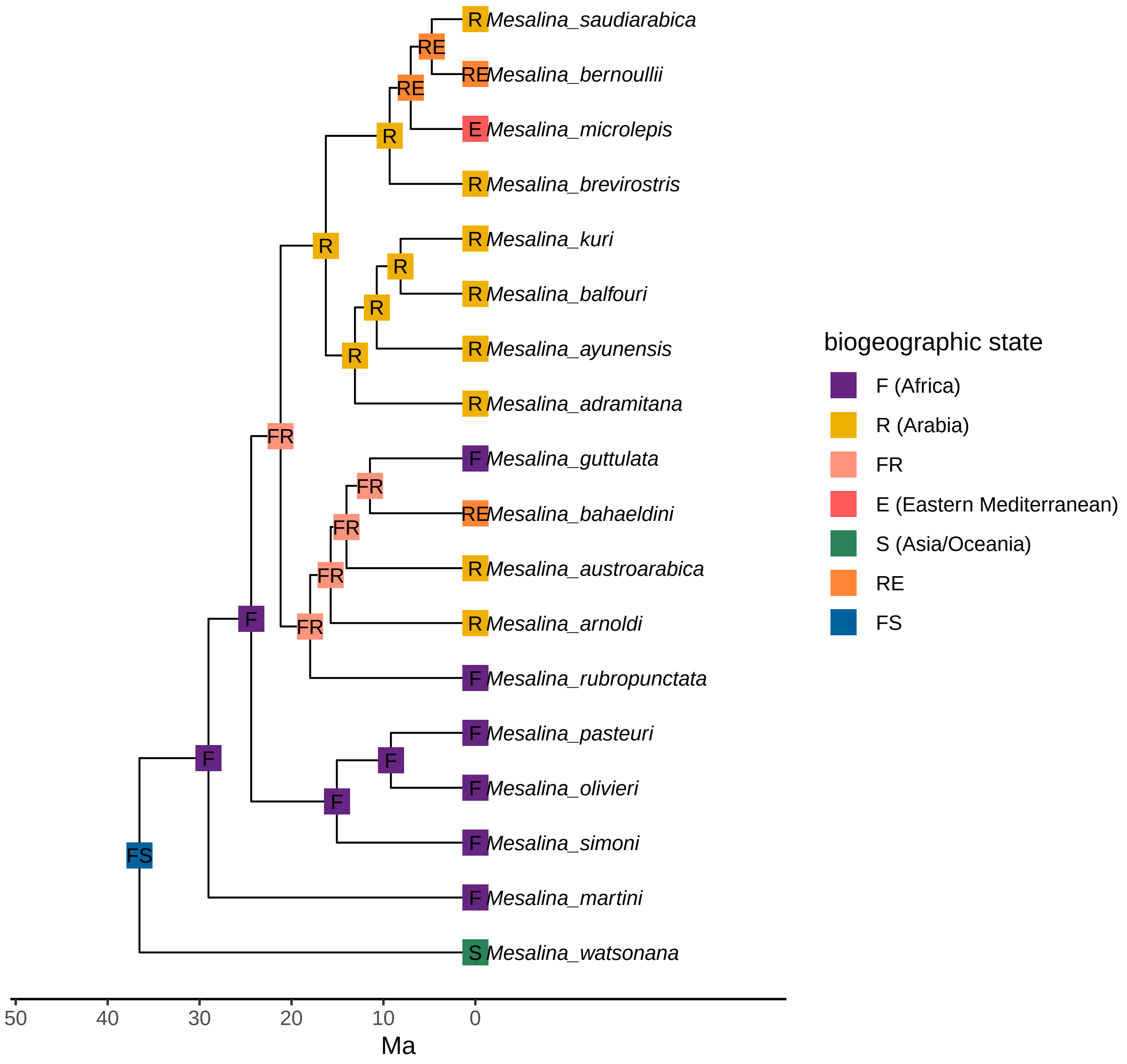
R Mesalina_saudiarabica
RE
RE Mesalina_bernoullii
RE
E Mesalina_microlepis
R
R Mesalina_brevirostris
R
R Mesalina_kuri
R
R Mesalina_balfouri
R
R Mesalina_ayunensis
R
R Mesalina_adramitana
FR
F Mesalina_guttulata
FR
RE Mesalina_bahaeldini
FR
R Mesalina_austroarabica
FR
R Mesalina_arnoldi
F
FR
F Mesalina_rubropunctata
F Mesalina_pasteuri
F
F
F Mesalina_olivieri
F
F Mesalina_simoni
FS
F Mesalina_martini
S Mesalina_watsonana
50
40
30
20
10
0
Ma
biogeographic state
F (Africa)
R (Arabia)
FR
E (Eastern Mediterranean)
S (Asia/Oceania)
RE
FS

*Naja*

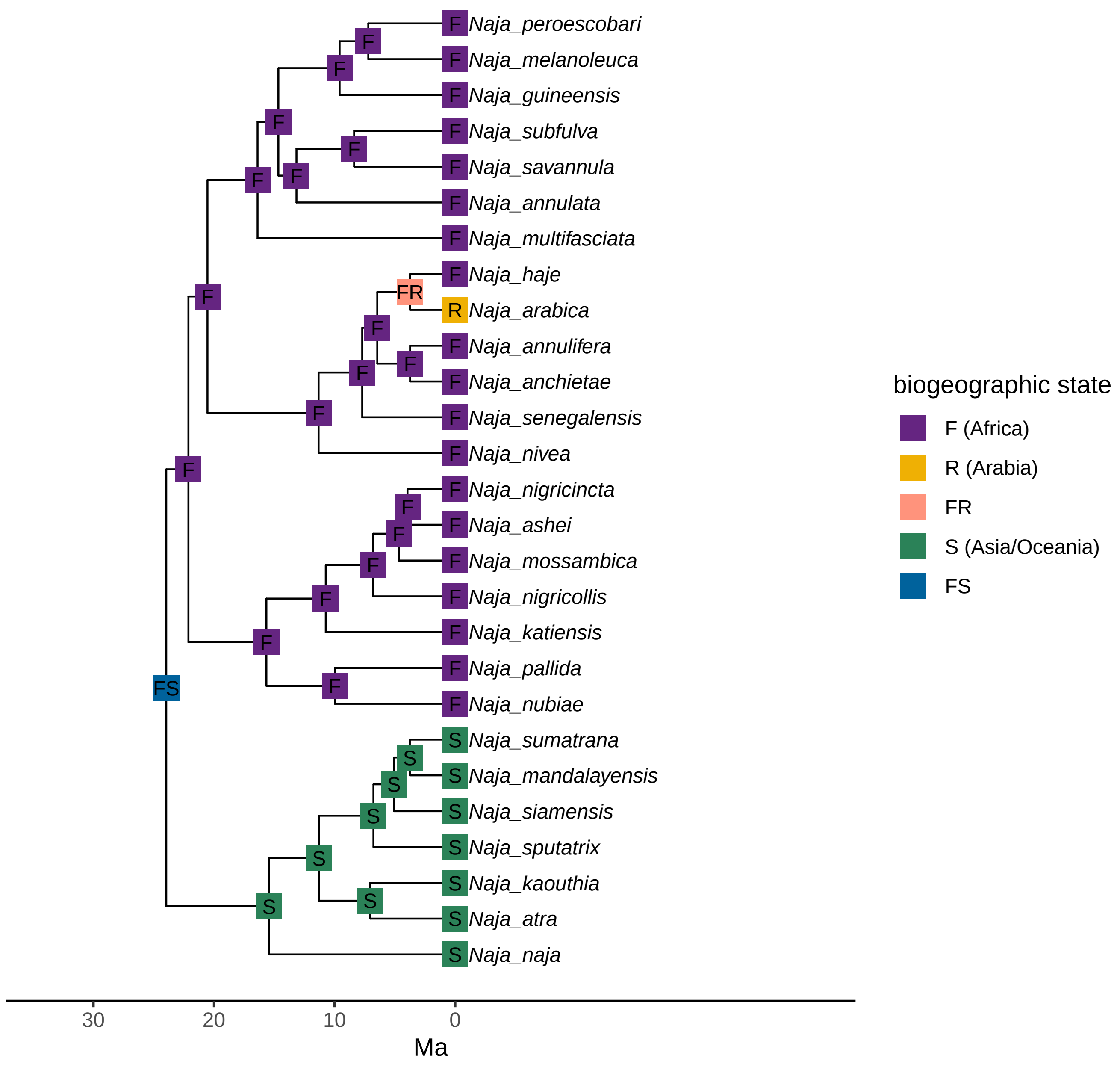
Naja_peroescobari
Naja_melanoleuca
Naja_guineensis
Naja_subfulva
Naja_savannula
Naja_annulata
Naja_multifasciata
Naja_haje
Naja_arabica
Naja_annulifera
Naja_anchietae
Naja_senegalensis
Naja_nivea
Naja_nigricincta
Naja_ashei
Naja_mossambica
Naja_nigricollis
Naja_katiensis
Naja_pallida
Naja_nubiae
Naja_sumatrana
Naja_mandalayensis
Naja_siamensis
Naja_sputatrix
Naja_kaouthia
Naja_atra
Naja_naja
FR
FS
30
20
10
0
Ma
biogeographic state
F (Africa)
R (Arabia)
FR
S (Asia/Oceania)
FS

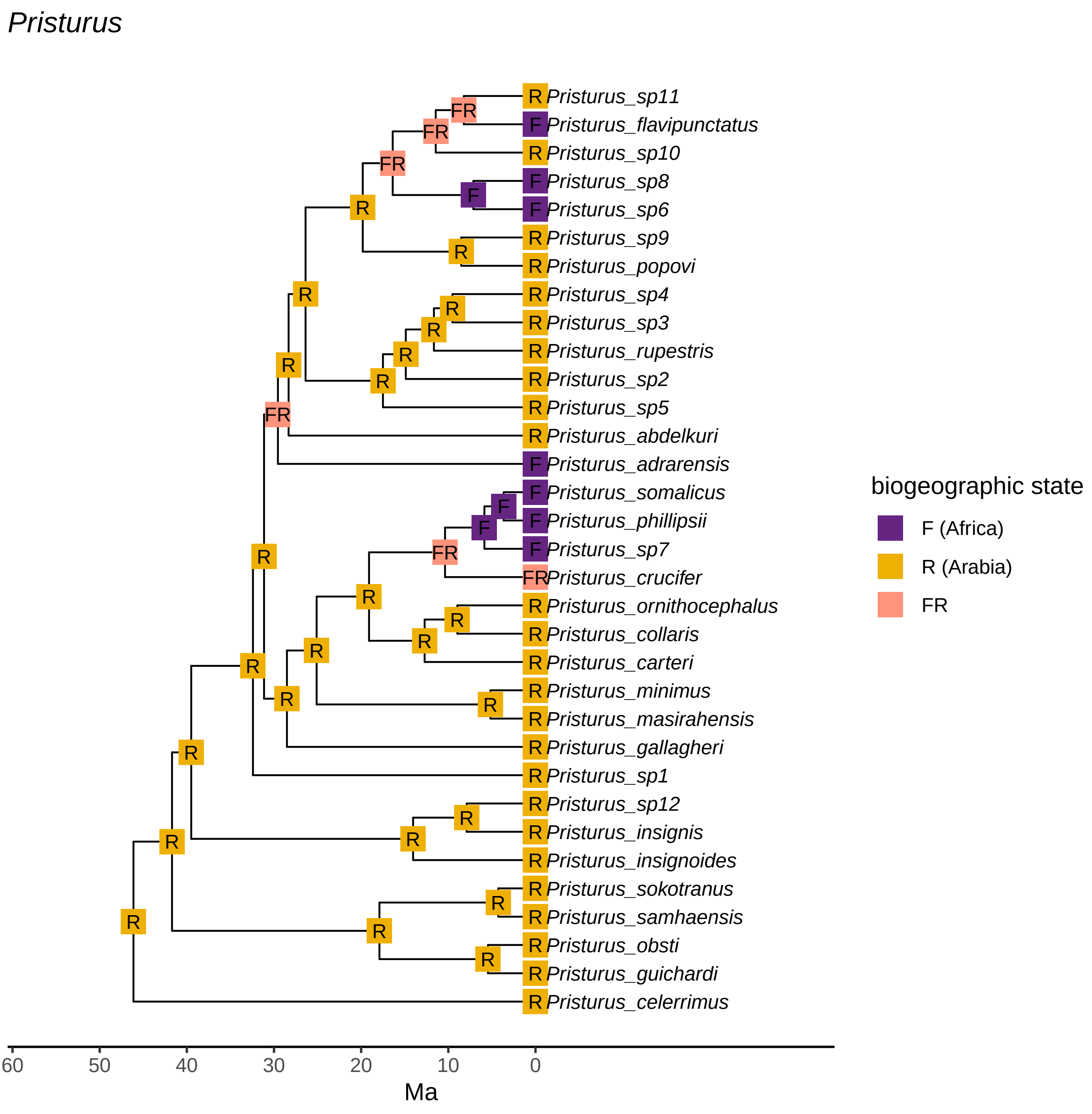
Pristurus
R Pristurus_sp11
F Pristurus_flavipunctatus
R Pristurus_sp10
F Pristurus_sp8
F Pristurus_sp6
R Pristurus_sp9
R Pristurus_popovi
R Pristurus_sp4
R Pristurus_sp3
R Pristurus_rupestris
R Pristurus_sp2
R Pristurus_sp5
R Pristurus_abdelkuri
F Pristurus_adrarensis
F Pristurus_somalicus
F Pristurus_phillipsii
F Pristurus_sp7
FR Pristurus_crucifer
R Pristurus_ornithocephalus
R Pristurus_collaris
R Pristurus_carteri
R Pristurus_minimus
R Pristurus_masirahensis
R Pristurus_gallagheri
R Pristurus_sp1
R Pristurus_sp12
R Pristurus_insignis
R Pristurus_insignoides
R Pristurus_sokotranus
R Pristurus_samhaensis
R Pristurus_obsti
R Pristurus_guichardi
R Pristurus_celerrimus
60
50
40
30
20
10
0
Ma
biogeographic state
F (Africa)
R (Arabia)
FR

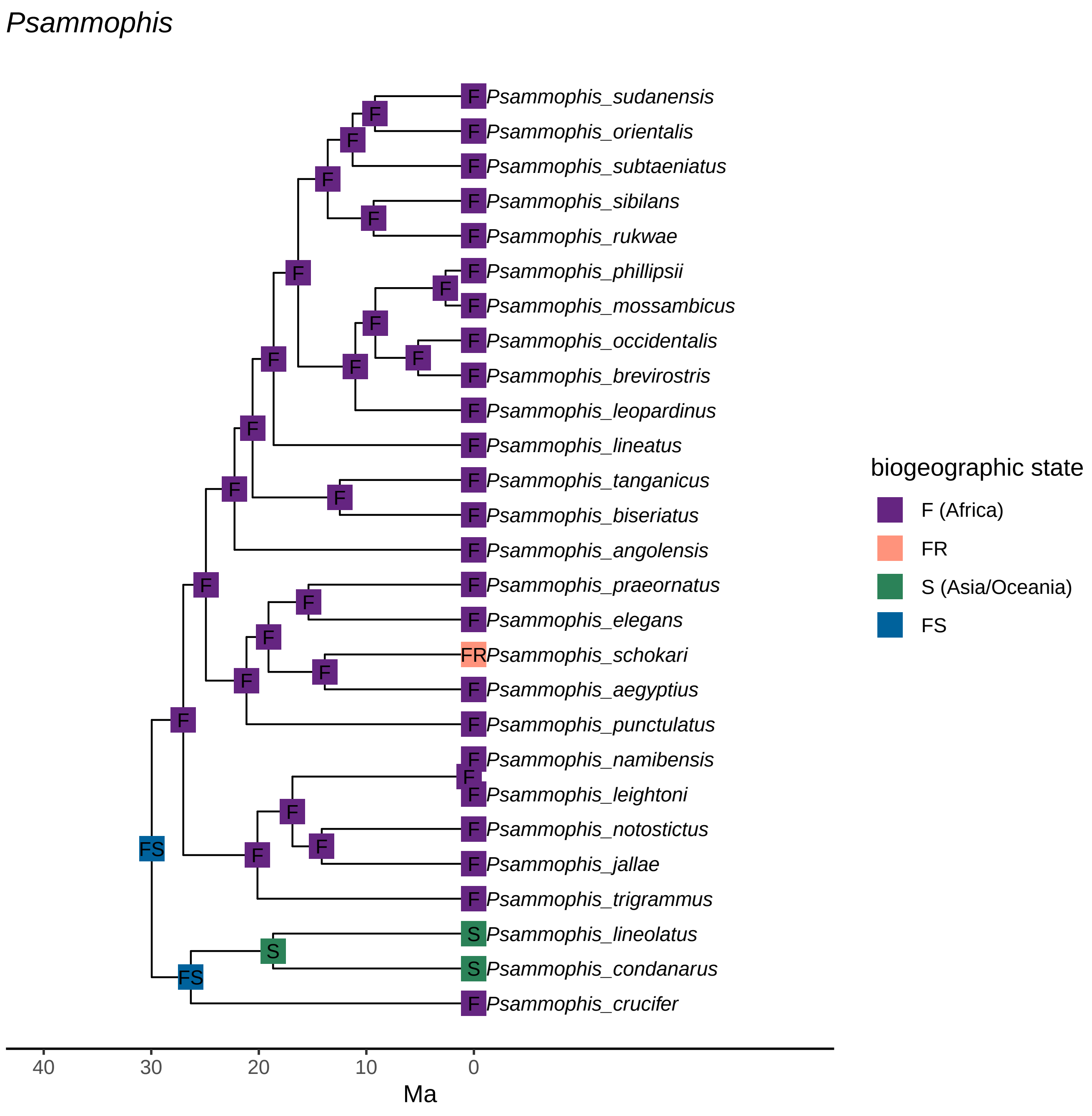
Psammophis
Psammophis_sudanensis
Psammophis_orientalis
Psammophis_subtaeniatus
Psammophis_sibilans
Psammophis_rukwae
Psammophis_phillipsii
Psammophis_mossambicus
Psammophis_occidentalis
Psammophis_brevirostris
Psammophis_leopardinus
Psammophis_lineatus
Psammophis_tanganicus
Psammophis_biseriatus
Psammophis_angolensis
Psammophis_praeornatus
Psammophis_elegans
Psammophis_schokari
Psammophis_aegyptius
Psammophis_punctulatus
Psammophis_namibensis
Psammophis_leightoni
Psammophis_notostictus
Psammophis_jallae
Psammophis_trigrammus
Psammophis_lineolatus
Psammophis_condanarus
Psammophis_crucifer
biogeographic state
F (Africa)
FR
S (Asia/Oceania)
FS
40
30
20
10
0
Ma

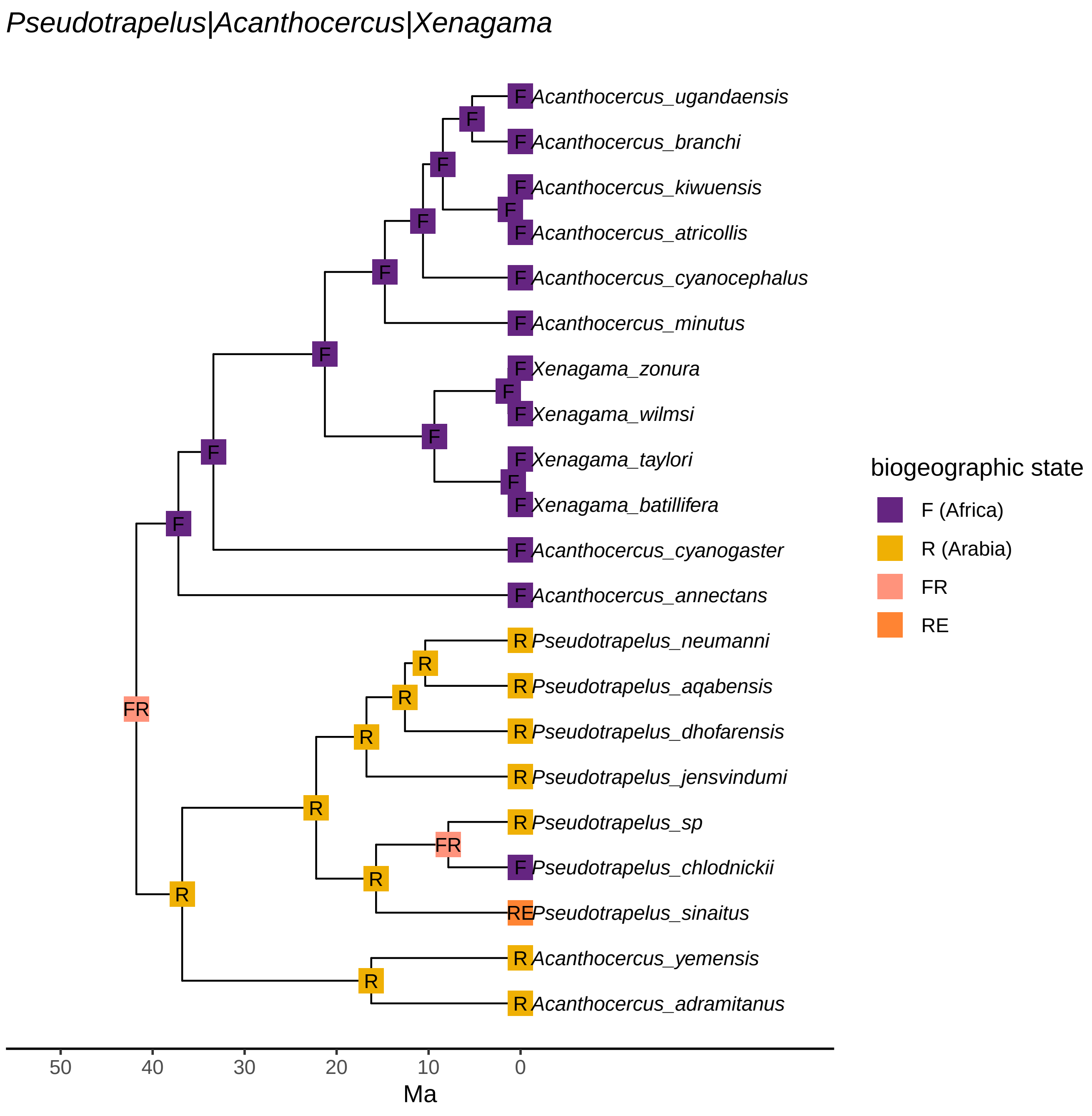

Pseudotrapelus|Acanthocercus|Xenagama
Acanthocercus_ugandaensis
Acanthocercus_branchi
Acanthocercus_kiwuensis
Acanthocercus_atricollis
Acanthocercus_cyanocephalus
Acanthocercus_minutus
Xenagama_zonura
Xenagama_wilmsi
Xenagama_taylori
Xenagama_batillifera
Acanthocercus_cyanogaster
Acanthocercus_annectans
Pseudotrapelus_neumanni
Pseudotrapelus_aqabensis
Pseudotrapelus_dhofarensis
Pseudotrapelus_jensvindumi
Pseudotrapelus_sp
Pseudotrapelus_chlodnickii
Pseudotrapelus_sinaitus
Acanthocercus_yemensis
Acanthocercus_adramitanus
biogeographic state
F (Africa)
R (Arabia)
FR
RE
50
40
30
20
10
0
Ma

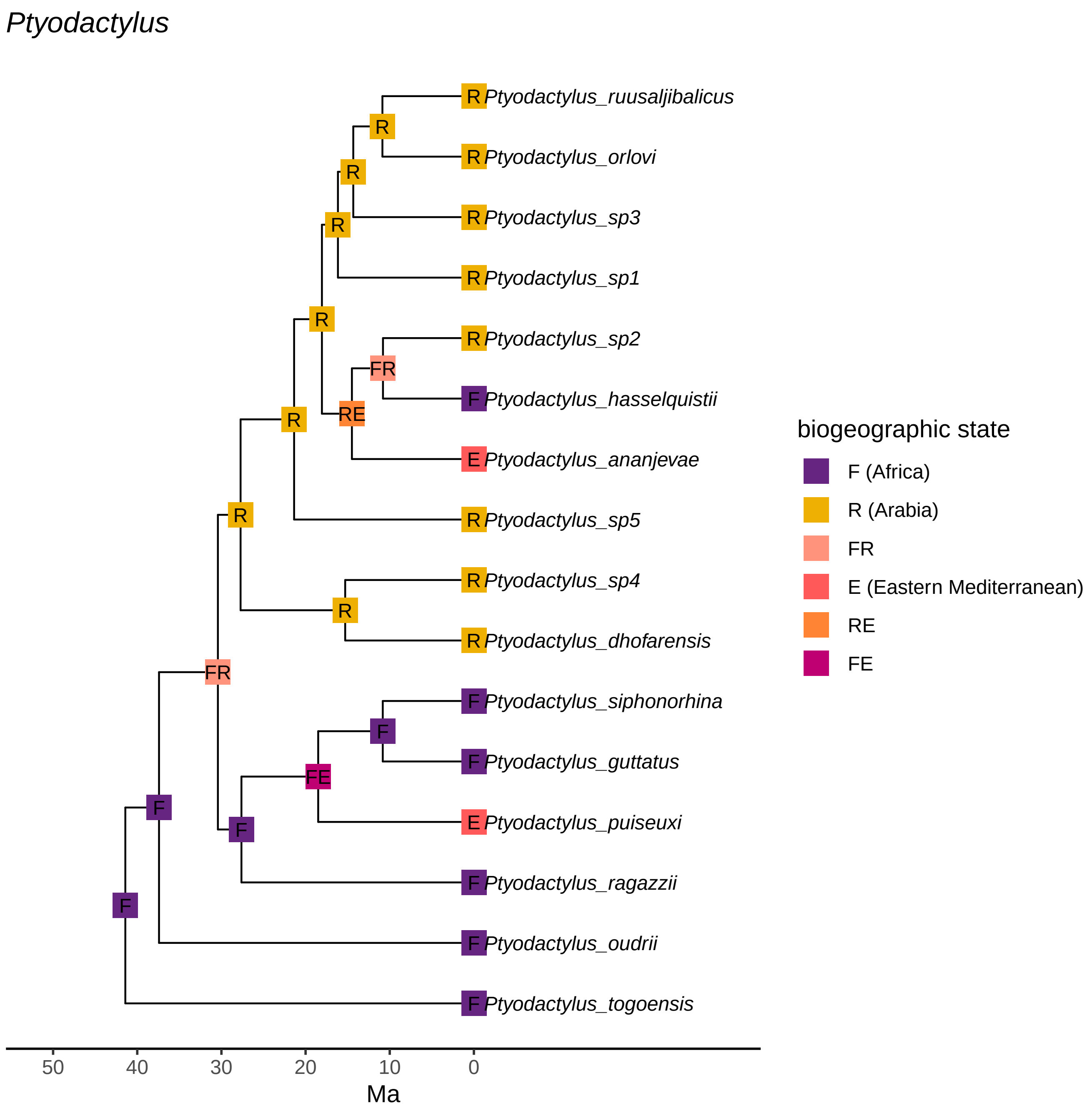
Ptyodactylus
R Ptyodactylus_ruusaljibalicus
R Ptyodactylus_orlovi
R Ptyodactylus_sp3
R Ptyodactylus_sp1
R Ptyodactylus_sp2
F Ptyodactylus_hasselquistii
E Ptyodactylus_ananjevae
R Ptyodactylus_sp5
R Ptyodactylus_sp4
R Ptyodactylus_dhofarensis
F Ptyodactylus_siphonorhina
F Ptyodactylus_guttatus
E Ptyodactylus_puiseuxi
F Ptyodactylus_ragazzii
F Ptyodactylus_oudrii
F Ptyodactylus_togoensis
R
R
R
R
FR
RE
R
R
R
FR
F
FE
F
F
F
50
40
30
20
10
0
Ma
biogeographic state
F (Africa)
R (Arabia)
FR
E (Eastern Mediterranean)
RE
FE

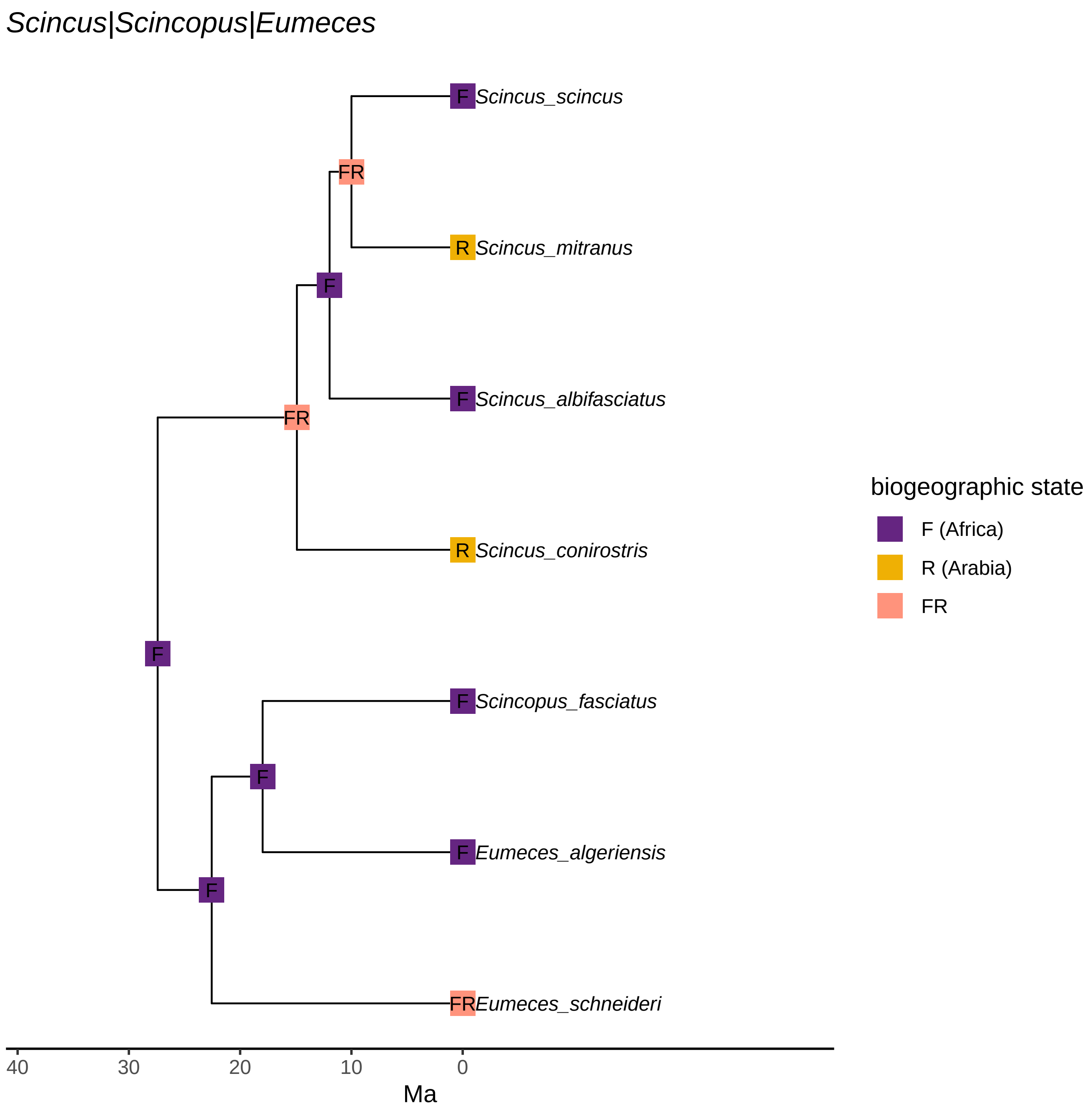
Scincus|Scincopus|Eumeces
F Scincus_scincus
FR
R Scincus_mitranus
F
F Scincus_albifasciatus
FR
R Scincus_conirostris
F
F Scincopus_fasciatus
F
F Eumeces_algeriensis
F
FR Eumeces_schneideri
40
30
20
10
0
Ma
biogeographic state
F (Africa)
R (Arabia)
FR

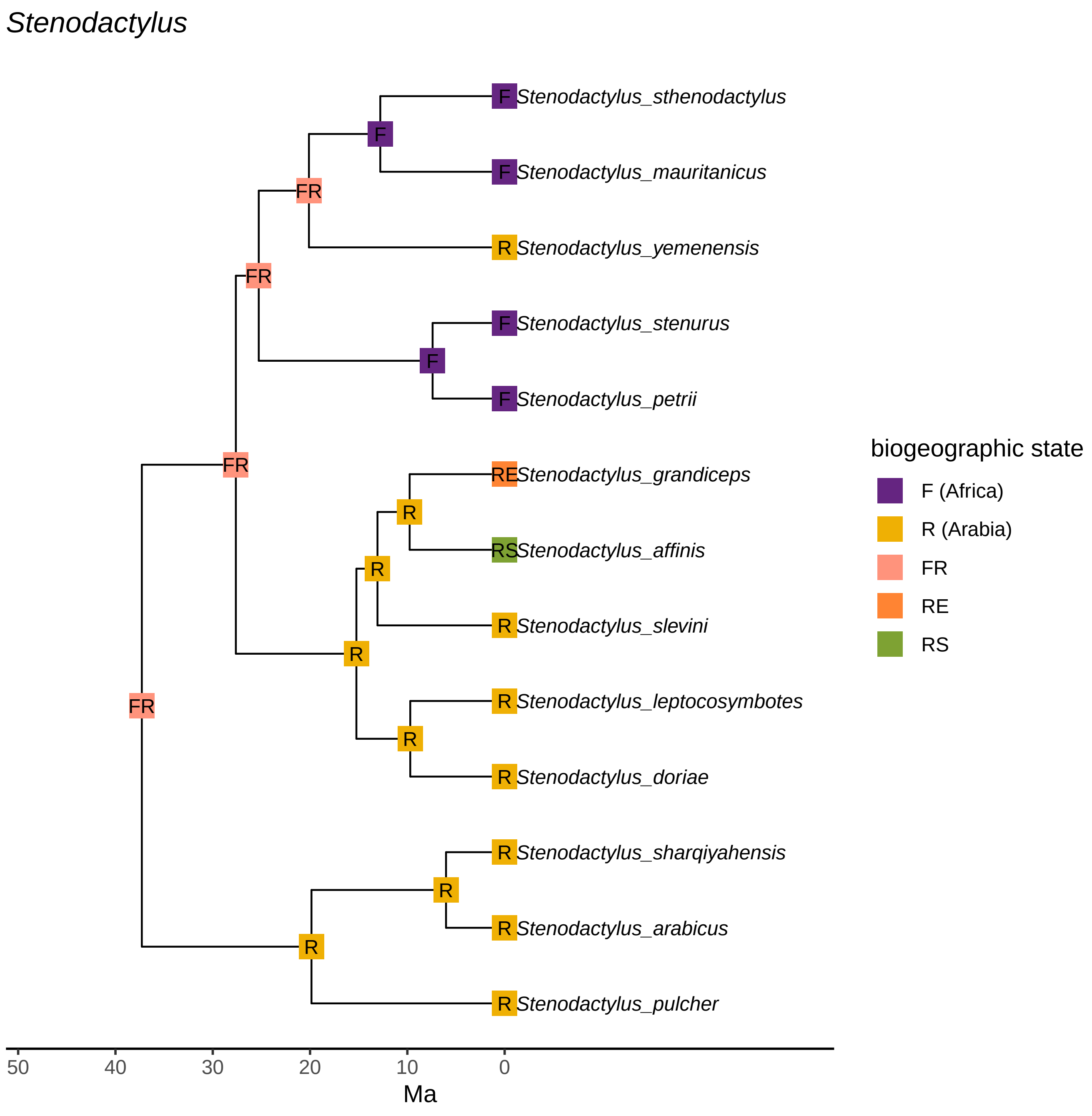
Stenodactylus
F Stenodactylus_sthenodactylus
F Stenodactylus_mauritanicus
R Stenodactylus_yemenensis
F Stenodactylus_stenurus
F Stenodactylus_petrii
RE Stenodactylus_grandiceps
RS Stenodactylus_affinis
R Stenodactylus_slevini
R Stenodactylus_leptocosymbotes
R Stenodactylus_doriae
R Stenodactylus_sharqiyahensis
R Stenodactylus_arabicus
R Stenodactylus_pulcher
F
FR
FR
F
FR
R
R
R
R
FR
R
R
50
40
30
20
10
0
Ma
biogeographic state
F (Africa)
R (Arabia)
FR
RE
RS

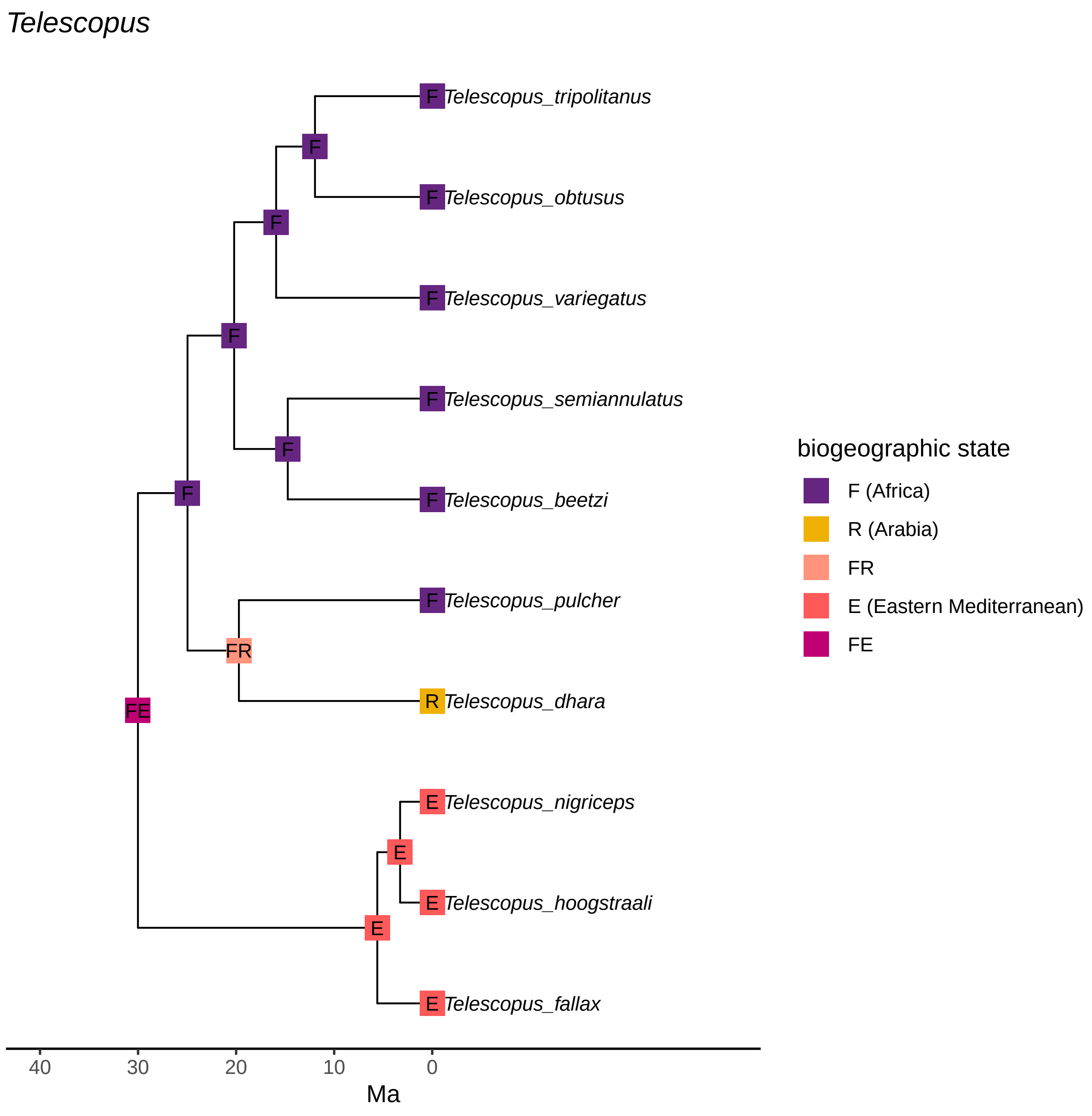
Telescopus
F
Telescopus_tripolitanus
Telescopus_obtusus
Telescopus_variegatus
Telescopus_semiannulatus
Telescopus_beetzi
Telescopus_pulcher
FR
R
Telescopus_dhara
FE
E
Telescopus_nigriceps
Telescopus_hoogstraali
Telescopus_fallax
40
30
20
10
0
Ma
biogeographic state
F (Africa)
R (Arabia)
FR
E (Eastern Mediterranean)
FE

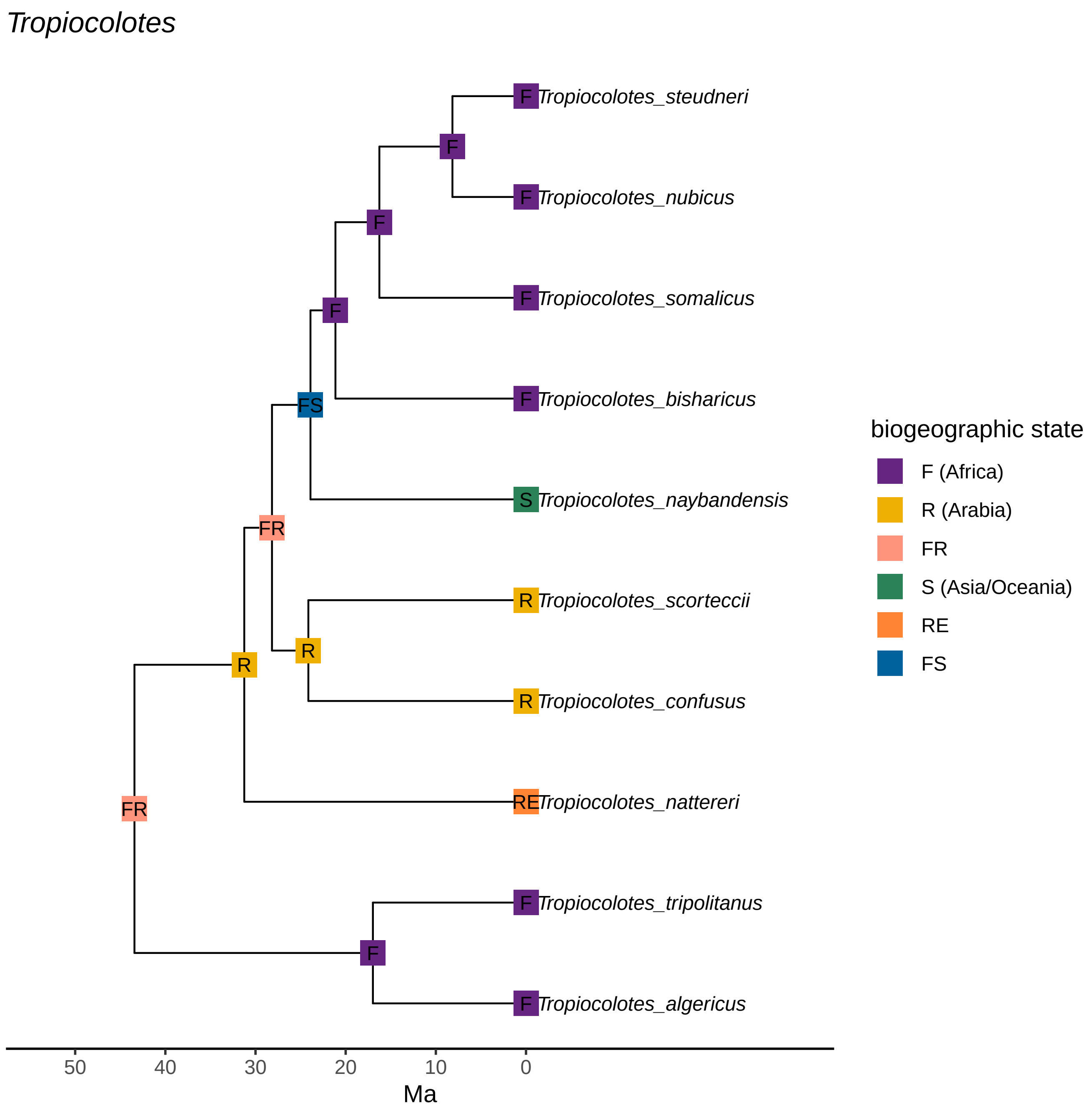
Tropiocolotes
Tropiocolotes_steudneri
Tropiocolotes_nubicus
Tropiocolotes_somalicus
Tropiocolotes_bisharicus
Tropiocolotes_naybandensis
Tropiocolotes_scorteccii
Tropiocolotes_confusus
Tropiocolotes_nattereri
Tropiocolotes_tripolitanus
Tropiocolotes_algericus
F
R
FR
FS
S
RE
50
40
30
20
10
0
Ma
biogeographic state
F (Africa)
R (Arabia)
FR
S (Asia/Oceania)
RE
FS

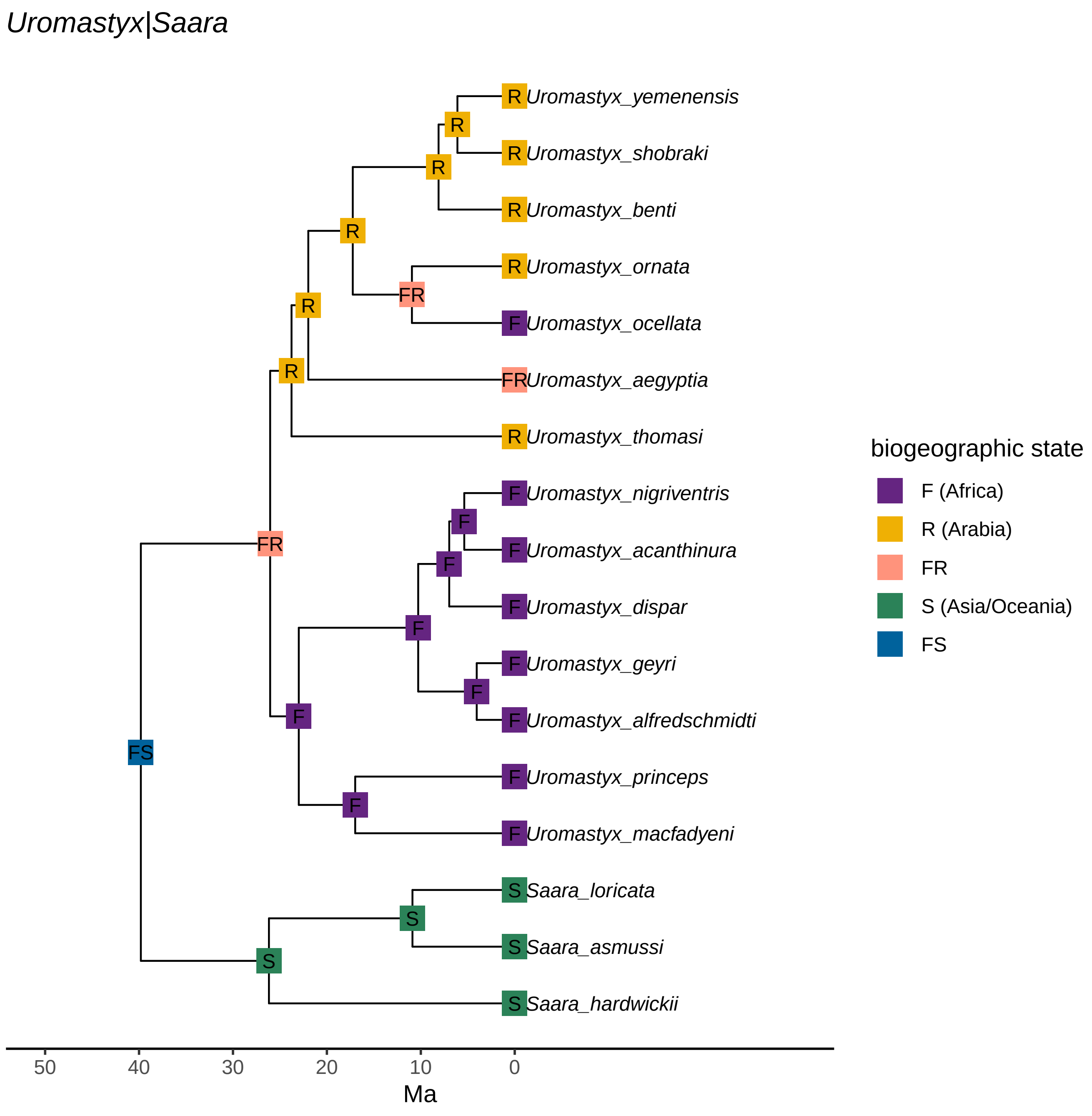
Uromastyx|Saara
R Uromastyx_yemenensis
R Uromastyx_shobraki
R Uromastyx_benti
R Uromastyx_ornata
F Uromastyx_ocellata
FR Uromastyx_aegyptia
R Uromastyx_thomasi
F Uromastyx_nigriventris
F Uromastyx_acanthinura
F Uromastyx_dispar
F Uromastyx_geyri
F Uromastyx_alfredschmidti
F Uromastyx_princeps
F Uromastyx_macfadyeni
S Saara_loricata
S Saara_asmussi
S Saara_hardwickii
FS
FR
50
40
30
20
10
0
Ma
biogeographic state
F (Africa)
R (Arabia)
FR
S (Asia/Oceania)
FS

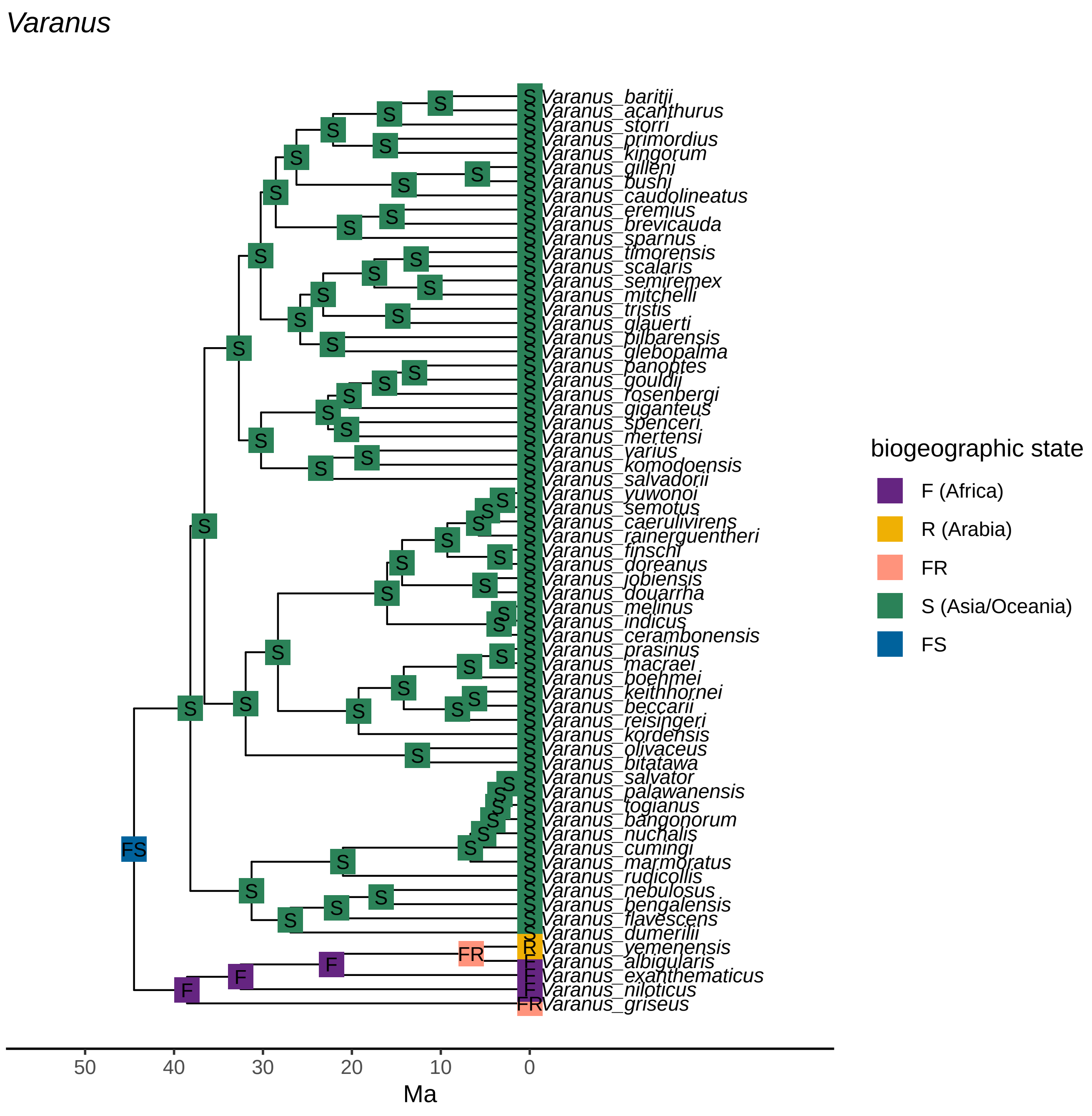
Varanus
Varanus_baritji
Varanus_acanthurus
Varanus_storri
Varanus_primordius
Varanus_kingorum
Varanus_gilleni
Varanus_bushi
Varanus_caudolineatus
Varanus_eremius
Varanus_brevicauda
Varanus_sparnus
Varanus_timorensis
Varanus_scalaris
Varanus_semiremex
Varanus_mitchelli
Varanus_tristis
Varanus_glauerti
Varanus_pilbarensis
Varanus_glebopalma
Varanus_panoptes
Varanus_gouldii
Varanus_rosenbergi
Varanus_giganteus
Varanus_spenceri
Varanus_mertensi
Varanus_varius
Varanus_komodoensis
Varanus_salvadorii
Varanus_yuwonoi
Varanus_semotus
Varanus_caerulivirens
Varanus_rainerguentheri
Varanus_finschi
Varanus_doreanus
Varanus_jobiensis
Varanus_douarrha
Varanus_melinus
Varanus_indicus
Varanus_cerambonensis
Varanus_prasinus
Varanus_macraei
Varanus_boehmei
Varanus_keithhornei
Varanus_beccarii
Varanus_reisingeri
Varanus_kordensis
Varanus_olivaceus
Varanus_bitatawa
Varanus_salvator
Varanus_palawanensis
Varanus_togianus
Varanus_bangonorum
Varanus_nuchalis
Varanus_cumingi
Varanus_marmoratus
Varanus_rudicollis
Varanus_nebulosus
Varanus_bengalensis
Varanus_flavescens
Varanus_dumerilii
Varanus_yemenensis
Varanus_albigularis
Varanus_exanthematicus
Varanus_niloticus
Varanus_griseus
50
40
30
20
10
0
Ma
biogeographic state
F (Africa)
R (Arabia)
FR
S (Asia/Oceania)
FS
